# Mitochondria dysfunction in Charcot Marie Tooth 2B Peripheral Sensory Neuropathy

**DOI:** 10.1101/2021.07.28.454213

**Authors:** Yingli Gu, Flora Guerra, Mingzheng Hu, Alexander Pope, Kijung Sung, Wanlin Yang, Simone Jetha, Thomas A. Shoff, Tessanya Gunatilake, Owen Dahlkamp, Linda Zhixia Shi, Fiore Manganelli, Maria Nolano, Yue Zhou, Jianqing Ding, Cecilia Bucci, Chengbiao Wu

## Abstract

Recent evidence has uncovered an important role of Rab7 in regulating mitochondrial morphology and function. Missense mutation(s) of Rab7 underlies the pathogenesis of Charcot Marie Tooth 2B (CMT2B) peripheral neuropathy. Herein, we investigated how mitochondrial morphology and function were impacted by the CMT2B associated Rab7^V162M^ mutation in fibroblasts from human CMT2B patients as well as in a knockin mouse model. In contrast to recently published results from studies of using heterologous overexpression systems, our results have demonstrated significant mitochondrial fragmentation in fibroblasts of both human CMT2B patients and CMT2B mouse embryonic fibroblasts (MEFs). Furthermore, we have shown that mitochondria were fragmented and axonal mitochondrial movement was dysregulated in primary cultured E18 dorsal root ganglion (DRG) sensory neurons, but not in E18 hippocampal and cortical primary neurons. We also show that inhibitors to either the mitochondrial fission protein Drp1 or to the nucleotide binding to Rab7 normalized the mitochondrial deficits in both MEFs and E18 cultured DRG neurons. Our study has revealed, for the first time, that expression of CMT2B Rab7 mutation at physiological level enhances Drp1 activity to promote mitochondrial fission, that may potentially underlie selective vulnerability of peripheral sensory neurons in CMT2B pathogenesis.

## Introduction

Mitochondria play an essential role in many facets of cellular functions: cellular energetics, metabolism, cell viability and cell death^1–3^. Mitochondria are extremely dynamic and able to divide, fuse, and move along the microtubule tracks to ensure their cellular distribution^3–5^. Mitochondrial dynamics allow the incessant and essential changes in mitochondrial size, shape and position inside the cells as well as their turnover^6^. In this context, an efficiently functioning mitochondrial network is assured by balanced fusion and fission processes.

In neurons, the mitochondrion is an essential organelle for metabolism and calcium homeostasis to maintain neuronal function and integrity^7, 8^. Mitochondrial dysfunction and altered mitochondrial dynamics are observed in a wide range of conditions, from impaired neuronal development to various neurodegenerative diseases^1, 2, 9–14^. Indeed, mitochondrial impairment in trafficking and function and alterations in mitochondrial turnover, which includes organelle biogenesis and quality check, are clearly associated with neurological disorders^2–4, 6, 9, 10, 15^. Neuronal cells are characterized by a complex morphology and functional sophistication linked to the remarkable length of their processes and to the requirement of rapid metabolic changes. These cells are particularly dependent on mitochondrial functions, on fission/fusion equilibrium and on mitochondrial localization^16–22^. Although the rules governing these changes and their functional significance are not fully understood, dysfunction of mitochondrial dynamics has been identified as a pathogenetic factor for disorders of both the central and the peripheral nervous systems^23^.

In recent years, several mutations in genes encoding proteins involved in regulation of mitochondrial dynamics and function have been identified in patients with Charcot-Marie-Tooth (CMT) peripheral neuropathy. CMT is characterized by different clinical, physiological, pathological and genetic phenotypes caused by alteration in more than 80 genes^24–28^, most of which encode proteins involved in the regulation of membrane traffic^29–31^.

The CMT type 2B (CMT2B) is a dominant, axonal form of the disease, characterized by sensory loss, progressive distal weakness, reduced tendon reflexes and normal or near-normal nerve conduction velocities. CMT2B is also an ulcero-mutilating neuropathy for foot deformities and high frequency of ulcers and infections leading to toe and foot amputations ^29, 32, 33^. CMT2B is caused by 5 missense mutations (p.L129F, p.K157N, p.N161T/I and p.V162M) in the *RAB7A* gene, encoding a GTPase of the RAB family^34–37^. Recently, another RAB7A mutation (p.K126R) characterized by a novel sensorimotor CMT2B phenotype was described^38^.

RAB7A, distinct from RAB7B, and hereafter referred to as RAB7, is ubiquitously expressed and has a pivotal role in the regulation of late endocytic trafficking^39, 40^. RAB7 also regulates apoptosis, membrane channel trafficking, and retromer recruitment^41–44^. In addition, RAB7 controls autophagosome maturation^45–49^. Notably, RAB7 has also specific roles in neurons as it regulates controlling neurotrophin trafficking and signaling, neurite outgrowth and neuronal migration during development^50–54^.

Although the biochemical and functional properties of the CMT2B-causing RAB7 mutant proteins have been previously investigated^54–58^, the exact mechanism by which mutated RAB7, albeit ubiquitous, causes dysfunction in peripheral neurons is still not clear. Emerging evidence has suggested a key role of the crosstalk between mitochondria and lysosomes in cellular physiology and its dysregulation in neurodegenerative disease as mitochondrial impairment can influence lysosomal function and *vice versa*^59^. Rab7 has been shown to regulate mitochondrial structure, motility and function and it was co-immuno-precipitated with the mitochondrial fusion protein MFN2^11^; Additionally, Rab7 was found to regulate phosphorylation of dynamin-related protein 1 (Drp1)^60^. Increased phosphorylation of Drp1 (pS616) promotes mitochondrial fission^61–63^. In particular, it has been demonstrated that RAB7 GTP hydrolysis is essential to regulate the duration, frequency and number of lysosomes-mitochondria contact^64^. Moreover, prolonged inter-mitochondrial contacts and defective mitochondrial motility was described in multiple CMT2 disease-linked mutations such as MFN2 (CMT2A), RAB7 (CMT2B) and TRPV4 (CMT2C)^65^. Interestingly, it was shown that Rab7 is involved in the translation of mRNAs encoding mitochondrial proteins at the late endosomal level, and CMT2B-causing Rab7 mutations markedly decreased axonal protein synthesis, impaired mitochondrial function, and compromised axonal viability^66^. These studies have provided important insights into the potential pathogenic mechanisms by which Rab7 mutation(s) alters mitochondrial dynamics leading to peripheral sensory neuropathy in CMT2B.

RAB7 is also described as a regulator of mitophagy mechanism controlling expansion of the LC3-positive isolation membrane around damaged mitochondria during mitophagy^44, 67^. Moreover, its activity is controlled by TBC1D5, the retromer-associated RAB7-specific GAP, which interacts with the subunit Vps29 of the retromer^68^. Due to this interaction, RAB7A localizes around damaged mitochondria and promotes their removal through Parkin-mediated mitophagy^69^. In the context of mitochondrial quality check, RAB7 is also responsible of mitochondrial derived vesicles (MDV) fusion with multivesicular bodies (MVBs) functioning as mitochondrial antigen-presenting system in immune cells via MDV trafficking in the absence of PINK1 or Parkin^70^.

The observation that mitochondrial impairment affects lysosomal functions and *vice versa* further supports interrelationship between the two organelles^59^. For instance, lysosomal activity is impaired in the setting of deficient mitochondrial respiration and disruption of endolysosomal trafficking^71^. Along similar lines, depletion or inhibition of apoptosis-inducing factor (AIF), OPA1, or PINK1 in neurons impairs lysosomal activity, thereby inducing accrual of autophagic substrates^72^. In light of this premises, it is reasonable to suppose that Rab7 might represent the mediator of inter-organelle communication and its dysfunction affect their interplay inducing massive alterations in peripheral neurons.

In this work, for the first time, fibroblasts of CMT2B patient harboring Rab7^V162M^ as well as a Rab7^V162M^ knockin mouse model were used to investigate mitochondrial structure and function in this pathology. Our data clearly demonstrate significant mitochondrial fragmentation in human patient fibroblasts and CMT2B mouse primary embryonic fibroblasts (MEFs) and dysregulation of mitochondrial morphology and axonal transport in primary cultured dorsal root ganglion (DRG) neurons.

## Results

### Human fibroblasts from CMT2B Rab7^V162M^ patients show significant mitochondrial fragmentation

Previous studies have shown that mitochondria were elongated by transient overexpression of CMT2B Rab7 constructs^64, 66^. To test whether this is the case under conditions that CMT2B-Rab7 mutant was expressed at physiological level, we cultured fibroblasts from three CMT2B patients that were heterozygous for Rab7^V162M^, as well as from two age-matched healthy control donors. The cells were cultured and incubated with 50nM Mito Tracker at 37°C for 40 min, live cell imaging was carried out to capture mitochondria as described in the Materials and Methods. Overall, mitochondria in fibroblasts from healthy controls showed elaborated networks with many tubular structures (Fig. 1A, B). These features were especially apparent in the zoom-in insets (Fig. 1A, B) and individual mitochondrion often appeared elongated (Fig. 1A, B, insets). To our surprise, the mitochondrial network complexity in CMT2B patient fibroblasts was significantly reduced with a concomitant increase of smaller and shorter structures (Fig. 1C, D). In fact, mitochondria with tubular structures were hard to find as compared with healthy controls (Fig. 1C, D versus 1A, B). Significant mitochondrial fragmentation was further detected in the zoom-in insets (Fig. 1C, D).

**Figure 1.**
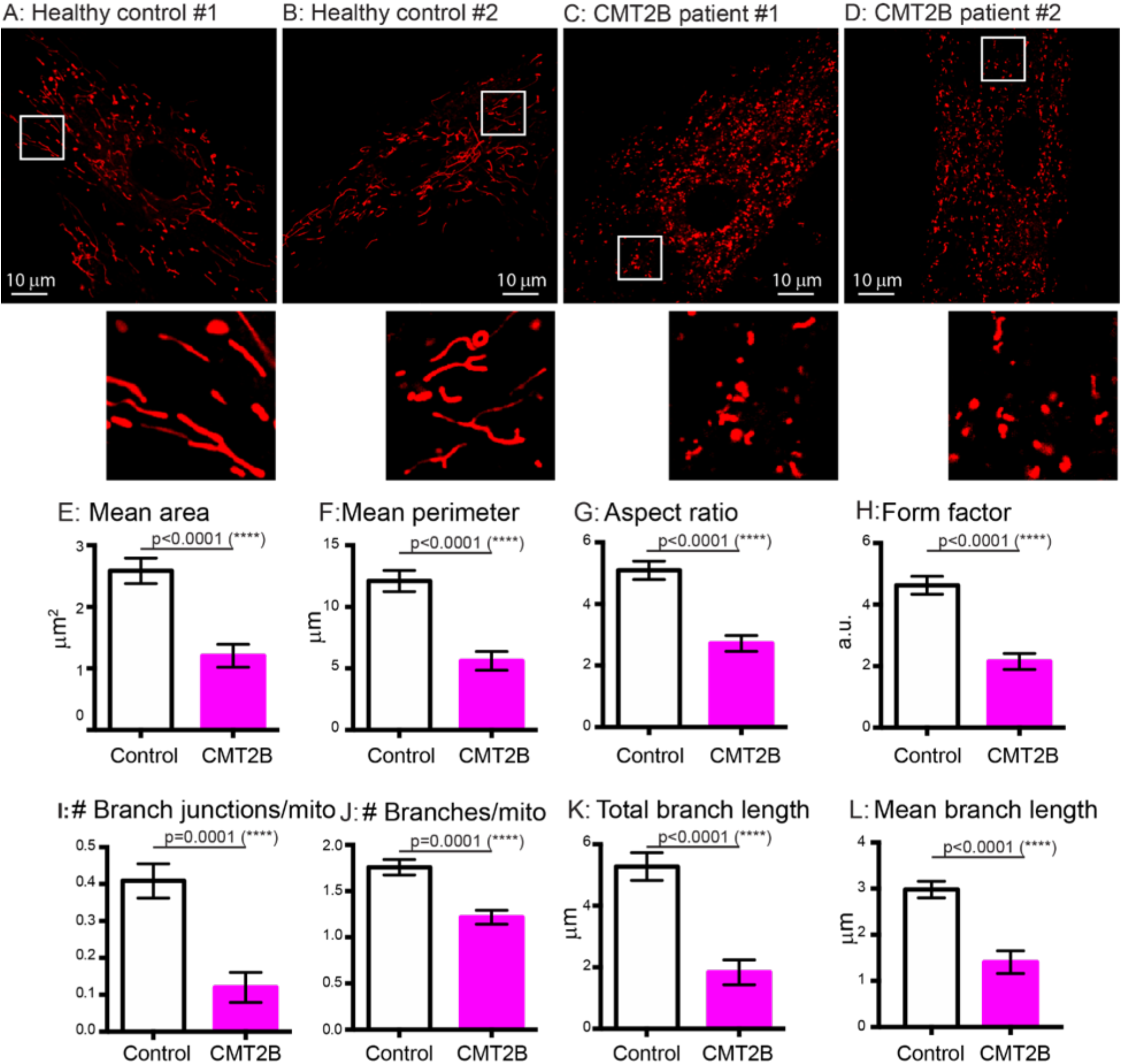
Mitochondria show significant fragmentation in skin fibroblasts from RAB7^V162M^ patients. Human skin fibroblasts from two healthy controls and two CMT2B patients were cultured as described in Materials and Methods. Fibroblasts were incubated with Mito Tracker and mitochondrial images were captured by live cell imaging. A, B: representative images of mitochondria from two healthy controls; C, D: representative images of mitochondria from two CMT2B patients. To better illustrate the mitochondria morphology, a small inset from each image (white box) was magnified and presented. The images were analyzed and quantitated using the Mitochondria Analyzer Plugin in Fiji (ImageJ). The measurements for mean area (E), mean perimeter (F), aspect ratio (G), form factor (H), the number of branch junctions/mitochondrion (I), the number of branches/mitochondrion (J), total branch length (K) and mean bran length (L) are presented. Results are shown as mean ± SEM. Significance analysis was carried out using Prism. Statistical significances were calculated by unpaired t-test. All p values are shown in the graphs.

To quantitate the changes in mitochondrial morphology in CMT2B patient fibroblasts, we processed the raw images using the Mitochondria Analyzer in Fiji/ImageJ as described in the Materials and Methods. Our analyses have revealed that mitochondria in CMT2B patients exhibited a significant reduction in the mean area (1.204 ± 0.186 µm^2^ versus 2.587 ± 0.208 µm^2^ for healthy controls) (Fig. 1E) and perimeter (5.608 ± 0.767 µm versus 12.1 ± 0.857 µm for healthy controls) (Fig. 1F). In addition, both AR, the aspect ratio (2.717 ± 0.258 versus 5.092 ± 0.3 for healthy controls) (Fig. 1G) and the form factor (2.153 ± 0.261 versus 4.626 ± 0.291 for healthy controls) (Fig. 1H) were reduced in CTM2B patient fibroblasts. Since branch junction of mitochondria indicates the network connectivity, which is critical for remodeling mitochondrial morphology and activity^5^, the numbers of branch junction per mitochondrion were also measured as described previously^73, 74^. Our quantitative results confirmed that mitochondria in CMT2B patient fibroblasts, on average, showed a significant reduction in the number of branch junctions (0.120 ± 0.041 versus 0.409 ± 0.046 for healthy controls) (Fig. 1I), the number of branches (1.217 ± 0.075 versus 1.760 ± 0.084 for healthy controls) (Fig. 1J), the total length of branches (1.835 ±

0.402 µm versus 5.273 ± 0.448 µm for healthy controls) (Fig. 1K) as well as the average branch length (1.407 ± 0.245 µm versus 2.979 ± 0.180 µm for healthy controls) (Fig. 1L). Thus mitochondrial network complexity is reduced in CMT2B cells. Taken together, these results provided strong evidence that mitochondria in CMT2B patient fibroblasts undergo significant mitochondrial fragmentation leading to a reduction in both the mitochondrial network complexity and the mitochondrial size. These are both surprising and interesting findings, given that recent studies using transient overexpression systems showed CMT2B Rab7 mutants induced mitochondrial elongation^64, 66^.

### Mitochondria are fragmented in mouse embryonic fibroblasts from CMT2B RAB7^V162M^ knockin model

We next wondered if our findings with respect to mitochondrial fragmentation in CMT2B also held true in a model system in which mutant Rab7 was expressed at physiological level. To this end, we turned to a CMT2B-Rab7^V162M^ knockin mouse model (C57BL6). By mutating 484G to A in the Rab7 gene located on Exon 5 (Supplemental Fig. 1A), a V162>M switch was achieved in Rab7. By homologous recombination, the mutated allele (a G to A switch at 484 of Exon 5, corresponding to V162>M) was introduced into the mouse genome. The selection marker, Neo^R^ was inserted into the mutated allele, in which LoxP sites were also included. Genotyping was performed using the PCR primer pair (Supplemental Fig. 1B). The wildtype allele (wt) gives rise to a 373 bp fragment while the mutant allele (Rab7^V162M:Neo^) produces a 490 bp fragment. We have obtained all three genotypes: wt (+/+), heterozygote (fln/+) and homozygote (fln/fln) with a typical Mendelian segregation ratio. Both the fln/+ and fln/fln pups survive to full adulthood (Supplemental Fig. 1C). Again, this mouse model will enable the study of fln/fln genotypes for possible gene dosage effects. Because a total knockout of Rab7 (-/-) in mice is embryonically lethal^75^, the fact that the fln/fln mice survived to the adulthood indicates that the Rab7^V162M^ mutation is unlikely to result in a total loss of function, a finding that is contradictory to a recent study of a fly model of CMT2B^76^. Therefore, fundamental differences do exist between the fly and mammalian mouse CMT2B models with regarding to Rab7 function. Studies with the mouse model will yield novel findings regarding the pathogenesis of CMT2B.

We cultured primary MEFs from E18 embryos of +/+, fln/+, fln/fln. Mitochondrial morphology in MEFs of 1-2 passages was analyzed by Mito Tracker labeling and live cell imaging as described for human fibroblasts. As shown in Fig. 2A, B and C, in MEFs of both fln/+ and fln/fln, mitochondria had significantly smaller area (0.890 ± 0.007µm^2^, 0.782 ± 0.005µm^2^, respectively) than +/+ mitochondria (1.014 ± 0.008µm^2^) (Fig. 2D). The changes in mitochondrial area were further revealed by the histogram distributions of mitochondrial area (Fig. 2D). Compared to +/+, the population of small size mitochondria (< 0.4 µm^2^) in the fln/+, fln/fln were increased; The population of medium size mitochondria (0.4-1.0 µm^2^) and the large size mitochondria (>1.0 µm^2^) were reduced, with the large size mitochondria showing a more significant reduction (Fig. 2D). Concomitantly, the mean perimeter of mitochondria, another indicator of mitochondrial size, had a very similar reduction in both fln/+ and fln/fln (5.189 ± 0.037 µm, 4.572 ± 0.025 µm, respectively) compared to +/+ mitochondria (6.017 ± 0.043 µm) (Fig. 2E). The changes in mitochondrial perimeter were also revealed by the corresponding histogram distributions (Fig. 2E). Compared to +/+, the population of small size mitochondria (< 2.5 µm) in the fln/+, fln/fln were increased; The medium size mitochondria (2.5-7.5 µm) were reduced with the large size mitochondria (>7.5 µm) showing a more significant reduction (Fig 2E). These data have demonstrated significantly reduced mitochondrial size in CMT2B RAB7^V162M^ MEFs.

**Figure 2.**
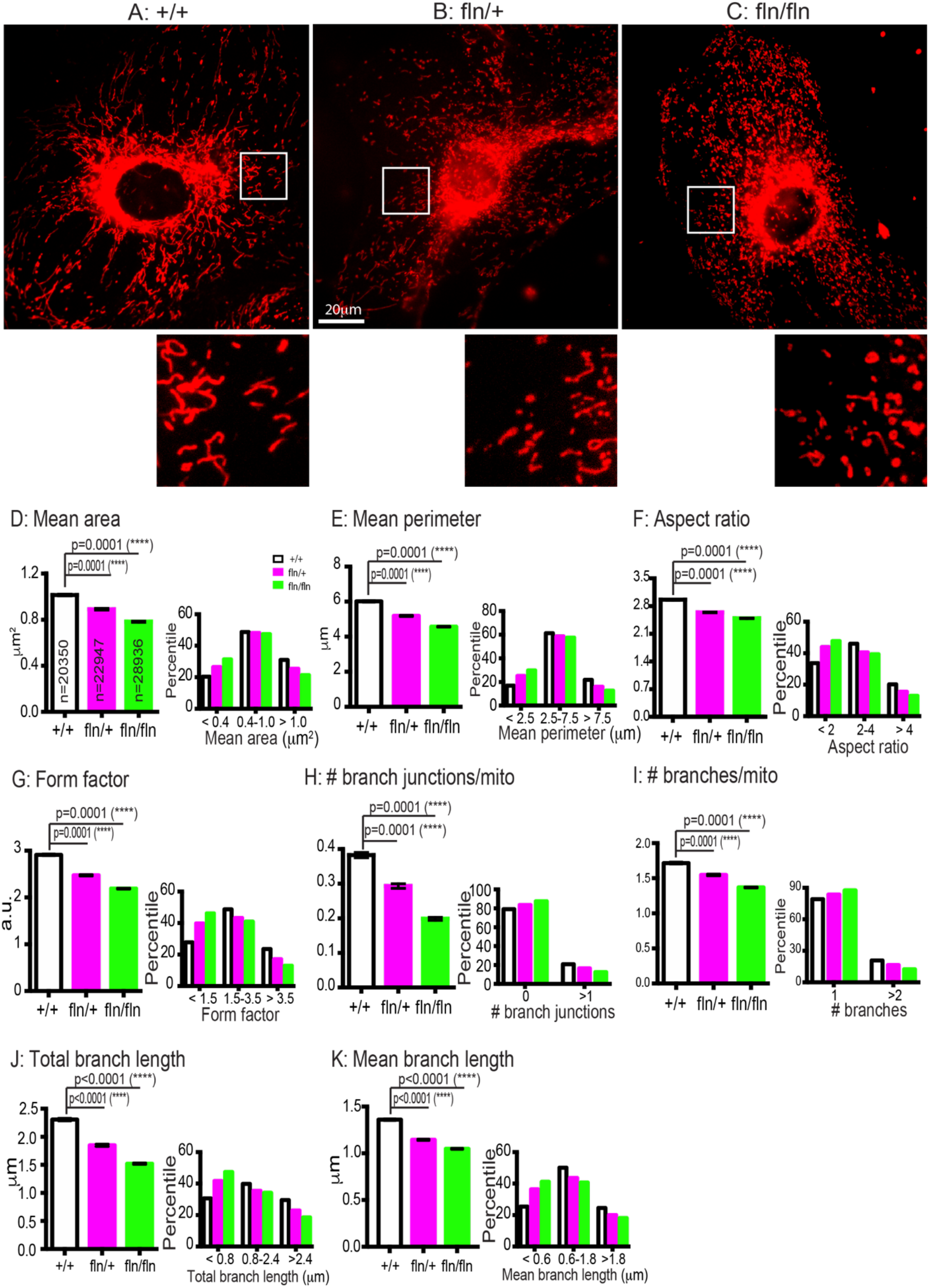
Mitochondria show significant fragmentation in MEFs from RAB7^V162M^ mutant mice. Mouse embryonic skin fibroblasts from +/+, fln/+, fln/fln E18 mice were dissected, cultured and maintained as described in Materials and Methods. MEFs were incubated with Mito Tracker and images were captured by live-cell imaging. A-C: representative images of mitochondria in +/+, fln/+, fln/fln. To better illustrate the mitochondria morphology, a small inset from each image (white box) was magnified and presented. The images were analyzed and quantitated using the Mitochondria Analyzer Plugin in Fiji (ImageJ). The measurements for mean area (D), mean perimeter (E), aspect ratio (F), form factor (G), the number of branch junctions/mitochondrion (H), the number of branches/mitochondrion (I), total branch length (J) and mean branch length (K) are presented. In addition, histogram distributions of mitochondria with respect to these measurements are also included in each perspective parameter. Results are shown as mean ± SEM. Significance analysis was carried out using Prism. Statistical significances were calculated by One-Way ANOVA. All *p* values are shown in the graphs.

The aspect ratio (AR) in fln/+ and fln/fln MEFs was measured at 2.640 ± 0.010 and 2.491 ± 0.008, respectively, significantly shorter than the +/+ MEF mitochondria (2.960 ± 0.012) (Fig. 2F). Likewise, histogram distributions of AR showed that short length (< 2) of mitochondria were increased, while medium (2-4) and long length (> 4) of mitochondria were reduced in both the fln/+, fln/fln MEFs as compared to +/+ MEFs (Fig. 2F). For the mitochondria with curvatures, the form factor in fln/+ and fln/fln MEFs was measured at 2.475 ± 0.016 and 2.191 ± 0.011, respectively, also significantly shorter than the +/+ MEF mitochondria (2.912 ± 0.019) (Fig. 2G). Histogram distributions of FF showed that short length (< 1.5) of mitochondria were increased, with an accompanying reduction of medium (1.5-3.5) and long length (> 3.5) mitochondria in both the fln/+, fln/fln MEFs (Fig. 2G). Further, the morphological changes of mitochondria were more pronounced in the fln/fln cells than the fln/+ cells (Fig. 2D-G), suggesting a gene-dosage dependent effect. These data are consistent with our human fibroblast results (Fig. 1). We thus conclude that expression of Rab7^V162M^ mutant allele at physiological conditions induces significant mitochondrial fragmentation.

We next measured the mitochondrial network complexity. As for the average numbers of branch junction per mitochondrion, the value for the +/+ cells was 0.383 ± 0.008. However, the fln/+ MEFs exhibited a significant reduction (0.293 ± 0.007), the reduction was even more pronounced in the fln/fln MEFs (0.199 ± 0.004) (Fig. 2H). This was confirmed by the quantification of mitochondria without or with at least one branch junction, which showed the reduction of mitochondria with one or multi-branch junctions in the mutant groups (Fig. 2H). Concomitantly, the average numbers of branches per mitochondrion in the fln/+, fln/fln MEFs exhibited a significant reduction (1.547 ± 0.012, 1.369 ± 0.008, respectively) compared to the +/+ cells (1.716 ± 0.015) (Fig. 2I), which was confirmed by the analysis of distribution, showing the reduction of mitochondria with multi branches (> 2) in the mutant groups (Fig. 2I). For the measurement of total branch length, it was 2.308 ± 0.023 µm in the +/+ cells, while the values in both the fln/+ and the fln/fln MEFs were significantly reduced (1.849 ± 0.020 µm, 1.521 ± 0.014 µm, respectively) (Fig. 2J). The corresponding histogram distribution showed that short length (< 0.8 µm) of mitochondria were increased, while medium (0.8-2.4 µm) and long length (> 2.4 µm) of mitochondria were reduced in both the fln/+, fln/fln MEFs as compared to +/+ MEFs (Fig. 2J). Likewise, the mean branch length in the fln/+, fln/fln MEFs exhibited consistent reduction (1.144 ± 0.007 µm, 1.048 ± 0.006 µm, respectively) compared to the +/+ cells (1.359 ± 0.008 µm) (Fig. 2K), which was confirmed by showing an increase of short length (< 0.6 µm) of mitochondria accompanying with a reduction of medium (0.6-1.8 µm) and long length (> 1.8 µm) of mitochondria in CMT2B RAB7 mutant (Fig. 2K). From Fig. 2H-K, we noticed that the changes of mitochondrial network complexity in the fln/fln group also displayed a gene-dosage dependent effect contrast to fln/+ group. Taken together, these data have demonstrated that knockin expression of Rab7^V162M^ in MEFs results in a significant reduction in the mitochondrial network complexity, a finding consistent with our human fibroblast study (Fig. 1).

### Excessive mitochondrial fragmentation in fln/fln MEFs is rescued by inhibition of Drp1 and Rab7

In mammalian cells, mitochondrial division is regulated by dynamin-related Protein 1, Drp1. Increased Drp1 activity leads to excessive mitochondrial fission^77, 78^. In a recent study, Rab7 was shown to regulate the Drp1 activity^60^. Moreover, Drp1 is involved in mitochondrial fission in CMT associated with GDAP1 mutation^4, 79, 80^. We thus postulated that excessive mitochondrial fragmentation in fln/+, fln/fln MEFs was a result of increased Drp1 activity when the Rab7^V162M^ allele was expressed. In addition, based on our previously published studies demonstrating that CMT2B-Rab7 mutants all show increased binding to GTP^54, 57^, we also hypothesized that competitive nucleotide binding inhibitors of Rab7 would also reduce mitochondrial fragmentation.

We first tested if increased Drp1 activity was responsible for excessive mitochondrial fragmentation in the CMT2B mutant MEFs. We decided to treat fln/fln MEFs with 50μM Mdivi- 1, an effective inhibitor for Drp1^81, 82^. In parallel experiments, the fln/fln MEFs were also treated with CID1067700, a competitive nucleotide binding inhibitor of Rab7^49, 83^. Cells were treated with the vehicle (0.1% DMSO), 50μM Mdivi-1, 1µM and 10µM CID1067700 Mitochondria were captured with Mito Tracker staining by live cell imaging as described above. Representative images were shown in Fig. 3A-D. Our quantitative analyses have revealed that mitochondria in fln/fln cells treated with the DMSO vehicle control had an average area of 0.719 ± 0.005µm^2^ (Fig. 3E), consistent with earlier results (Fig. 2D); The mitochondrial area was significantly increased in cells treated either with Mdivi-1 (1.035 ± 0.008µm^2^), or with CID1067700 either at 1µM (0.827 ± 0.006µm^2^) or at 10µM (0.853 ± 0.005µm^2^). The changes are reflected by an increase in the percentiles of medium size and large size of mitochondria accompanying with a concomitantly reduction in the small size mitochondria population (histogram in Fig. 3E). Likewise, Mdivi-1, CID1067700 also increased the perimeter of mitochondria (DMSO: 4.348 ± 0.025 µm; Mdivi-1: 6.136 ± 0.043 µm; 1 µM CID10607700: 4.875 ± 0.030 µm; 10 µM CID10607700: 5.215 ± 0.030 µm) (Fig. 3F). As expected, the aspect ratio (Fig. 3G) was significantly rescued by Mdivi-1, low and high dosage of CID1067700 (DMSO: 2.536 ± 0.0093; Mdivi-1: 3.117 ± 0.012; 1 µM CID10607700: 2.621 ± 0.010; 10 µM CID10607700: 2.884 ± 0.010). Consistently, the form factor was also increased by the drug treatments (DMSO: 2.155 ± 0.011; Mdivi-1: 2.973 ± 0.019; 1 µM CID10607700: 2.347 ± 0.013; 10 µM CID10607700: 2.602 ± 0.013) (Fig. 3H). These data are evidence that inhibition of Drp1 and Rab7 restored the deficits in mitochondrial size.

**Figure 3.**
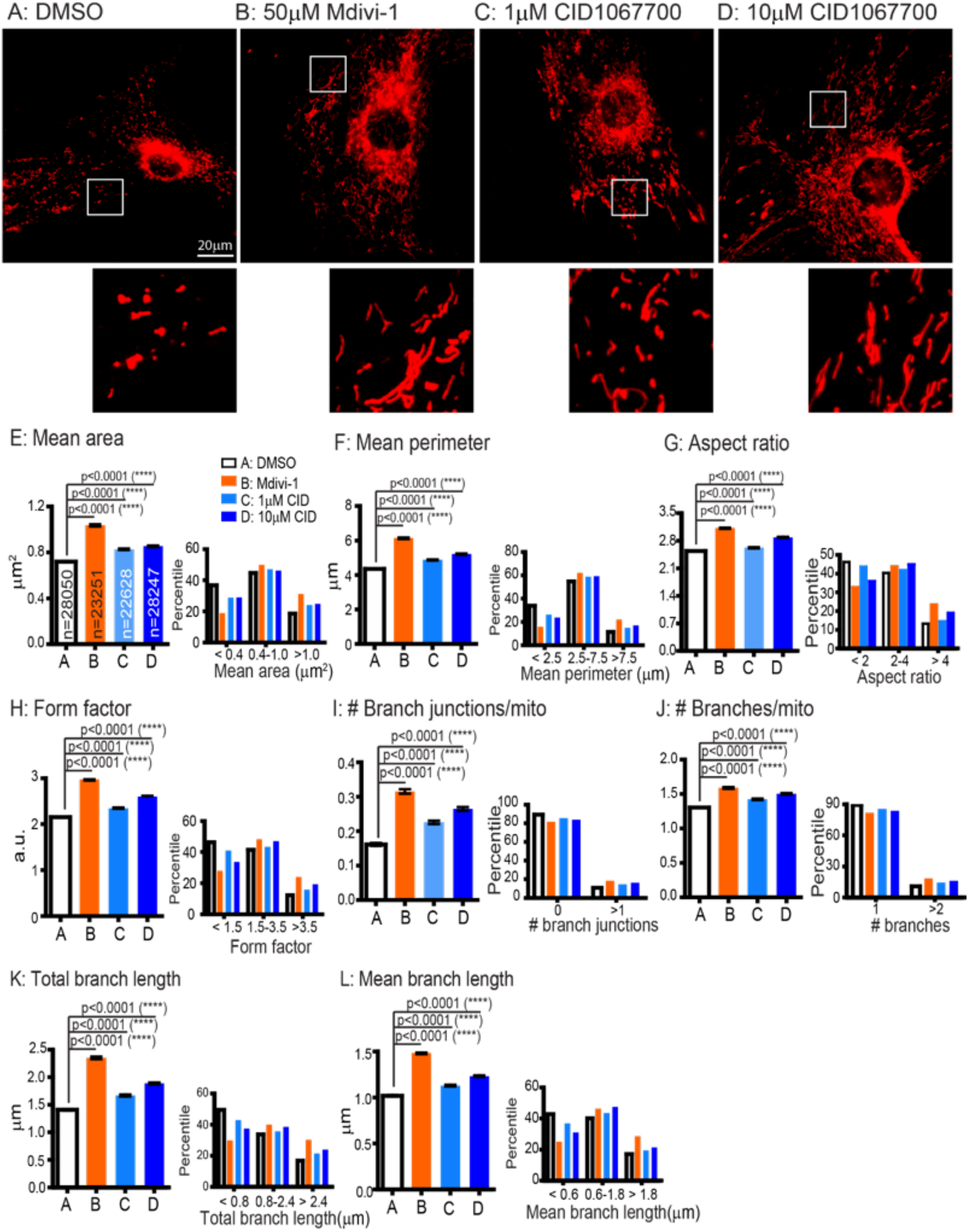
Mitochondrial fragmentation in fln/fln MEFs is rescued by both Mdivi-1 and CID1067700 treatment. MEFs from fln/fln E18 embryos were cultured and were treated DMSO, 50µM Mdivi-1, 1µM CID1067700, 10µM CID1067700. Mitochondria labeling, live cell imaging and mitochondrial quantitation were as in Fig. 2. A-D: Representative images for cells treated with DMSO, 50µM Mdivi-1, 1µM CID1067700, 10µM CID1067700. To better illustrate the mitochondria morphology, a small inset from each image (white box) was magnified and presented. The measurements for mean Area (D), mean perimeter (E), aspect ratio (F), form factor (G), the number of branch junctions/mitochondrion (H), the number of branches/mitochondrion (I), total branch length (J) and mean bran length (K) are presented. In addition, histogram distributions of mitochondria with respect to these measurements are also included in each perspective parameter. Results are shown as mean ± SEM. Significance analysis was carried out using Prism. Statistical significances were calculated by One-Way ANOVA. All *p* values are shown in the graphs.

Our results have also shown that the mitochondria network complexity was also rescued by treatment with Mdivi-1 and CID1067700; The number of branch junctions/mitochondrion was increased (DMSO control: 0.162 ± 0.004; Mdivi-1: 0.316 ± 0.006; 1 μM CID1067700: 0.226 ± 0.005; 10 μM CID1067700: 0.265 ± 0.005) (Fig. 3I); The treatments also resulted in an increase in the number of branches per mitochondrion (DMSO: 1.304 ± 0.007; Mdivi-1: 1.590 ± 0.012; 1 µM CID10607700: 1.423 ± 0.009; 10 µM CID10607700: 1.504 ± 0.009) (Fig. 3J). The branch length, including both the total branch (Fig. 3K) and mean branch length (Fig. 3L) of mitochondria was increased in response to the drug treatment (Total length: DMSO: 1.412 ± 0.013 µm; Mdivi-1: 2.352 ± 0.023 µm; 1 µM CID10607700: 1.669 ± 0.016 µm; 10 µM CID10607700: 1.888 ± 0.015 µm. Mean length: DMSO: 1.020 ± 0.006 µm; Mdivi-1: 1.484 ± 0.009 µm; 1 µM CID10607700: 1.129 ± 0.007 µm; 10 µM CID10607700: 1.232 ± 0.007 µm.). We thus conclude that inhibition of Drp1 activity and Rab7 nucleotide binding restores mitochondrial network in the fln/fln MEFs.

### Mitochondria are fragmented in axons of DRG sensory neurons in CMT2B mouse model

Pathologically, CMT2B is known to selectively afflict axonal functions of peripheral sensory neurons. To investigate whether or not peripheral sensory neurons had similar changes in mitochondrial morphology as MEFs in our CMT2B mouse model, we cultured DRG sensory neurons from +/+, fln/+ and fln/fln E18 embryos. We first attempted to analyze mitochondria in the soma of DRGs as we did in MEFs. Unfortunately, the average diameter of DRG neuronal soma at DIV5 of all three genotypes was about 10-20 μm (Fig. S2), comparing to ∼80-100 μm for either human fibroblasts (Fig. 1) or MEFs (Fig. 2). Under the same 100x magnification as we imaged MEFs, individual mitochondrion marked by Mito Tracker was extremely difficult to be discerned and to be quantitated in the soma of these DRG neurons (Fig. S2).

We then quantitated axonal mitochondria using Mito Tracker by live cell imaging. Unlike MEFs, the majority of mitochondria in axons was rod-shaped, mitochondria with branches were rarely seen in axons as shown in Fig. 4A. We thus measured the aspect ratio (AR). Mitochondria in +/+ had an AR value of 1.743 ± 0.070 (Fig. 4B); This value was significantly decreased in the fln/fln sample (1.485 ± 0.040, p=0.0035) (Fig. 4B). Even though the fln/+ samples also showed a decrease to 1.698 ± 0.062, the decrease did not reach statistical significance (Fig. 4B).

**Figure 4.**
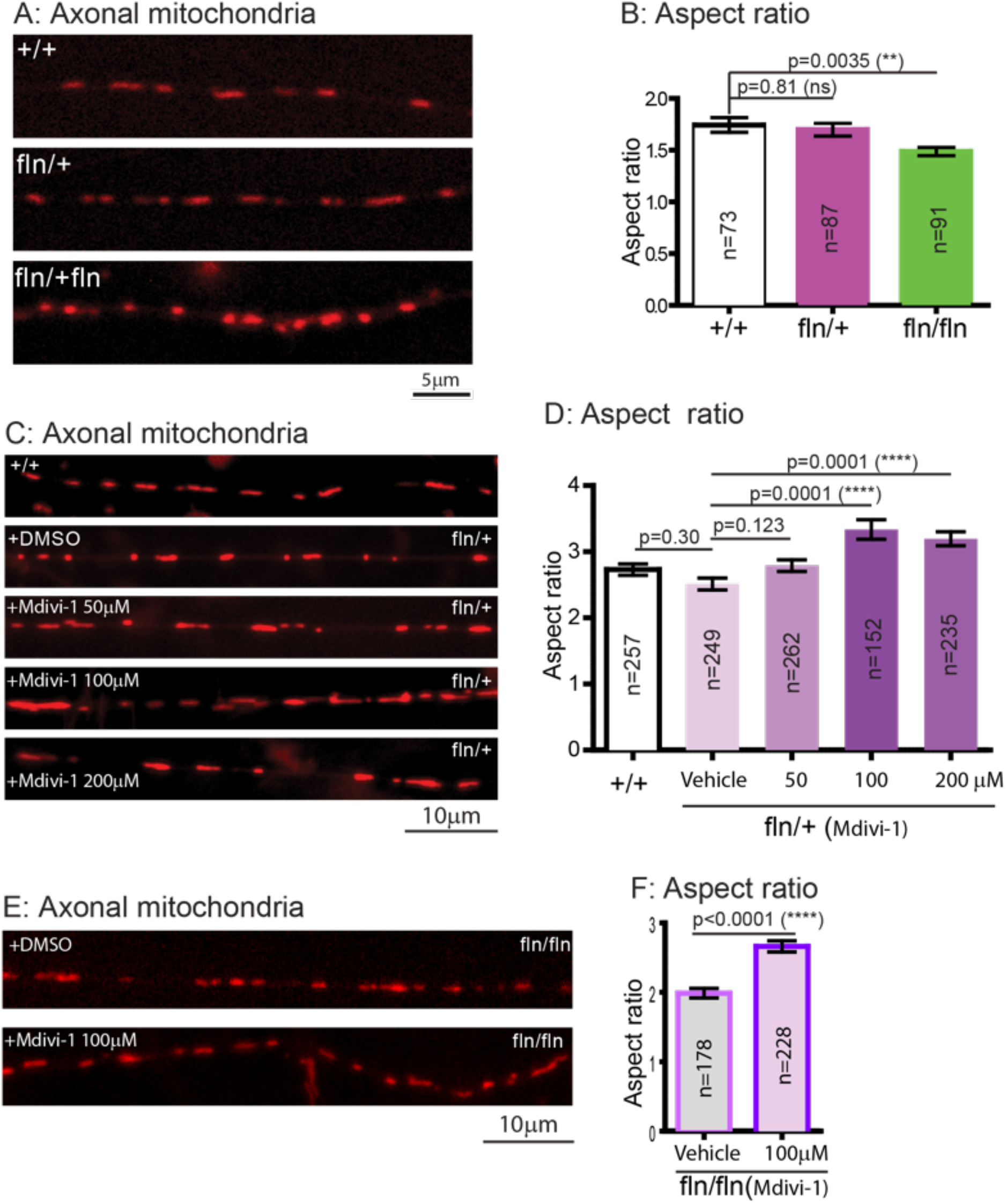
Mitochondria are significantly fragmented in axons of fln/fln E18 DRG sensory neurons of the CMT2B mutant mice. E18 DRG sensory neurons from +/+, fln/+, fln/fln embryos were dissected, cultured on PLL-coated cover-glasses and maintained as described in Materials and Methods. At DIV4-7, neurons were treated with Mito Tracker and time-lapsed images were captured with live imaging microscope. A: representative images of axonal mitochondria from +/+, fln/+, fln/fln; B: Aspect ratio for each group was measured as in Fig. 1-3. In C, representative images of mitochondria in axons of DRG neurons from untreated +/+, DMSO-treated fln/+ or fln/+ treated with 50 µM, 100 µM, 200 µM Mdivi-1. Quantitative measurements of aspect ratio under these conditions are presented in D. In E, representative images of mitochondria in axons of DRG neurons from fln/fln treated DMSO or 100 µM Mdivi-1. Quantitative measurements of aspect ratio under these conditions are presented in F. Results are shown as mean ± SEM. Significance analysis was carried out using Prism. Statistical significances were calculated by One-Way ANOVA or unpaired t-test. All *p* values are shown in the graphs.

To examine if Mdivi-1 could rescue axonal mitochondrial fragmentation in mutant DRG neurons, we first conducted a dose-dependent study using fln/+ DRG neurons. We treated the fln/+ DRG neurons with the DMSO vehicle, 50 µM, 100 µM, 200 µM of Mdivi-1 (final concentration) and measured the AR and compared these values to untreated +/+ DRG neurons (Fig. 4C). As expected, fln/+ DRG neurons treated with DMSO continued to show a decrease in AR, albeit the difference did not reach statistical significance (Fig. 4D); Treatment with Mdivi-1 increased the AR value and the increases were significant at 100 µM and 200 µM, the effect at 100 µM was maximal (2.817 ± 0.077 versus 2.260 ± 0.063 for DMSO, Fig. 4D). We next treated fln/fln DRG neurons with either DMSO or 100µM Mdivi-1 (Fig. 4E); Our quantitation revealed that Mdivi-1 significantly increased the AR value from 1.987 ± 0.070 in DMSO-treated samples to 2.661 ± 0.081 (Fig. 4F). These results have provided further evidence that mitochondrial morphology in DRG sensory neurons is impaired by the expression of CMT2B mutant allele and the effect is likely mediated by increased activity of Drp1.

### Mitochondria motility is increased in DRG sensory neurons of CMT2B mouse model

Previous studies have shown that one of main functions of CMT2B-causing RAB7A is to regulate long-range retrograde axonal transport in neurons^29, 50, 51, 53, 54, 84, 85^. We next examined axonal trafficking of mitochondria in DRG sensory neurons. To investigate this, the axonal movement of mitochondria in DRG cultures from three genotypes (+/+, fln/+, fln/fln) was labeled with Mito Tracker and time lapsed image series of axonal movement of mitochondria were recorded, as shown in Movie S1-S3. Kymographs were generated from the time lapsed image series and representative kymographs for each group were shown in Fig. 5A. We quantitated the average speed of mitochondrial transport in both the anterograde (Fig. 5B) and retrograde (Fig. 5C) direction. We also measured the percentile of moving mitochondria in the retrograde direction (Fig. 5D) as well as the percentile of total number of moving mitochondria (Fig. 5E). Interestingly, the average velocity of anterograde trafficking was slightly increased in fln/+ DRG neurons (1.324 ± 0.090 µm/s) and the increase in fln/fln DRG neurons reached statistical significance (1.388 ± 0.085 µm/s), when compared to that in the +/+ samples (1.094 ± 0.089 µm/s) (Fig. 5B). A similar trend was detected with respect to the average moving speed of mitochondria in the retrograde direction, i.e., a slight increase in the fln/+ (1.329 ± 0.093µm/s) and a significant increase in the fln/fln (1.659 ± 0.108 µm/s) samples when compared to 1.062 ± 0.101 µm/s in the +/+ DRG neurons (Fig. 5C). The percentile of retrograde trafficking mitochondria in the +/+ neurons was 40.5%, while this value was increased to 43.3%, 62.2% for the fln/+ and fln/fln DRG neurons, respectively (Fig. 5D). The percentile of total number of moving mitochondria in the +/+ neurons was 43.7%, this value was increased to 52.1% in the fln/+ and 55.9% in the fln/fln neurons (Fig. 4E).

**Figure 5.**
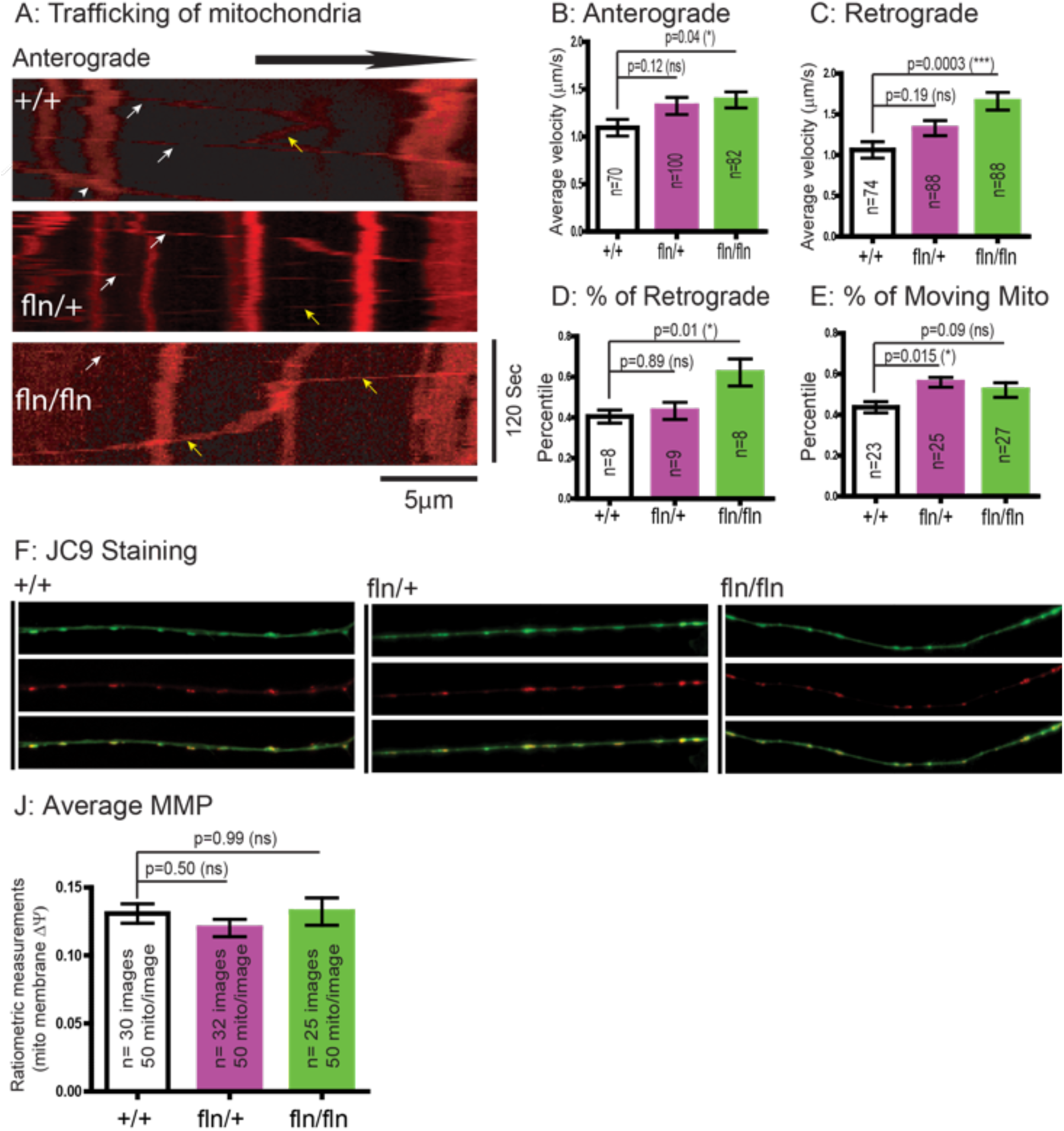
Axonal motility of mitochondria is altered in fln/fln E18 DRG sensory neurons of the CMT2B mutant mice. E18 DRG sensory neurons from +/+, fln/+, fln/fln embryos were dissected, cultured and labeled with Mito Tracker at DIV4-7 as in Fig. 4. Time-lapsed images of axonal mitochondria were captured by live imaging microscopy. The image series were used to generat kymographs in Image J to quantitate axonal mitochondrial motility. A: Representative kymographs of +/+, fln/+ and fln/fln. White arrows indicate trafficking in anterograde direction, yellow arrows indicate trafficking in retrograde direction. Average velocities of axonal mitochondria moving in the anterograde (B) and retrograde (C) direction. D: % of retrogradely moving mitochondria. E: % of total moving mitochondria. We also performed JC9 staining to measure MMP by ratiometric analysis. F: representative images of JC9 staining of mitochondria of DRG axons of +/+, fln/+, fln/fln. The green channel and the red channel images as well as the merged images are shown. J: Ratiometric measurements of MMP in +/+, fln/+, fln/fln. Results are shown as mean ± SEM. Significance analysis was carried out using Prism. Statistical significances were calculated by One-Way ANOVA. n.s.= non significance. All *p* values are shown in the graphs.

To measure any changes in mitochondrial membrane potential (MMP) in axonal mitochondria in DRG neurons, we next used stained +/+, fln/+, fln/fln DRG neurons with JC9, the ratiometric dye of mitochondria. Monomeric JC9 yields green fluorescence signal while increased MMP will result in aggregation of JC9 that will emit red fluorescence signal intensity. Representative images of green, red and merged images for each genotype were shown in Fig. 5F. To our surprise, we did not detect any significant changes in either the fln/+ or the fln/fln axonal mitochondria when compared to the +/+ axons (Fig. 5F). Our quantitation of the red/green ratio further confirmed our observation (Fig. 5G). Taken together, we conclude that mitochondrial motilities and moving speeds were increased in the mutant DRG neurons in a gene-dosage dependent manner, albeit no changes in MMP.

### Mitochondria are minimally affected in neurons of the central nervous system in the CMT2B mouse model

Given that Rab7 is ubiquitously expressed in all cell types ^40^ and clinically CMT2B patients are not known to suffer from any deficits in their brain functions. we next investigated if mitochondria were impacted in neurons from the central nerve system (CNS). We next cultured E18 hippocampal neurons and examined their mitochondria. Again, mitochondria in the soma of cultured E18 hippocampal neurons were difficult to measure due to the small diameter of these neurons (Fig. S3). We first performed JC9 staining to measure MMP. Representative images of green, red and merged images for +/+, fln/+, fln/fln were shown in Fig. 6A-C. Our ratiometric results have shown no significant changes in MMP in either fln/+ or fln/fln hippocampal neurons as compared to +/+ (Fig. 6D), which was confirmed by the histogram distribution (Fig. 6E). We next performed Mito Tracker labeling to measure the mitochondrial aspect ratio in the neurites. Our results have revealed no significant difference between the either fln/+ or fln/fln and +/+ in the shape and size of mitochondria (Fig. 6F), as measured by the aspect ratio (Fig. 6G). We further examined and quantitated mitochondrial transport in the neurites; Our results have shown that the fln/+ and fln/fln neurons did not show significant difference for the +/+ neurons in the average moving velocities, either in the anterograde (Fig. 6I) or in the retrograde direction (Fig. 6J). Interestingly, the percentile of retrograde moving mitochondria was significantly reduced in the fln/fln, but not in the fln/+, neurons when compared to that of +/+ (Fig. 6K). Yet, the percentile of moving mitochondria as a whole did not show any significant changes in either the fln/+ or fln/fln when compared to +/+ (Fig. 6L). Based on these findings, we have concluded that expression of the Rab7^V162M^ mutant allele in CMT2B knockin mice exhibits little impact on mitochondria motility and on membrane potential in E18 hippocampal neurons.

**Figure 6.**
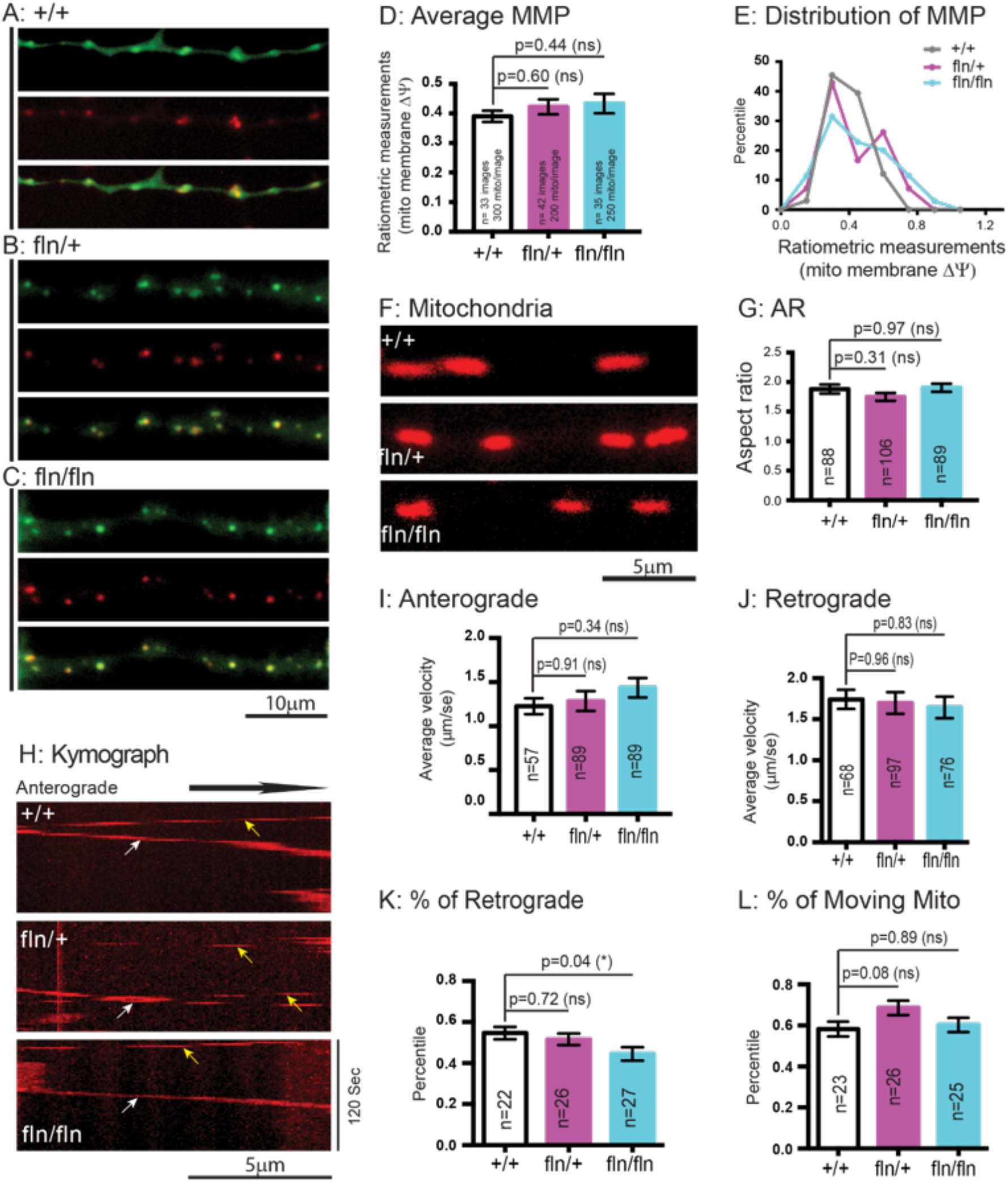
Analysis of mitochondria in E18 hippocampal neurons from CMT2B mutant mice. E18 hippocampal neurons from +/+, fln/+, fln/fln were dissected, cultured on PLL-coated cover- glasses and maintained as described in Materials and Methods. At DIV4-7, neurons were treated with JC9 and live-imaged. A-C: Representative images of mitochondria in neurites of E18 hippocampal neurons of +/+, fln/+ and fln/fln. D: Ratiometric measurements of MMP. E: Histogram analysis of MMP distribution. In F, Representative images of mitochondrial morphology in hippocampal neurons of +/+, fln/+, fln/fln. G: Aspect ratio measurements. Mitochondrial transport in neurites of hippocampal neurons of +/+, fln/+, flnfln was also captured by time lapsed imaging. The image series were used to generate kymographs and a representative image for each genotype is shown in H. White arrows indicate trafficking in anterograde direction, yellow arrows indicate trafficking in retrograde direction. Average velocities of axonal mitochondria moving in the anterograde (I) and retrograde (J) direction. K: % of retrogradely moving mitochondria. L: % of total moving mitochondria. Results are shown as mean ± SEM. Significance analysis was carried out using Prism. Statistical significances were calculated by One- Way ANOVA. n.s.= non significance. All *p* values are shown in the graphs.

To further test if Rab7^V162M^ mutant allele affects mitochondria in other populations of CNS neurons, we also cultured E18 cortical neurons from +/+, fln/+, fln/fln mouse embryos. Similar to cultured hippocampal neurons (Fig. S3), the soma size of cultured E18 cortical neurons was small making it difficult to measure the mitochondrial network in the soma of these neurons (Fig. S4.) As with the cortical neurons, we first performed JC9 staining and representative images of green, red and merged images for each genotype in neurites were shown in Fig. 7A-C. Our ratiometric analysis has revealed no significant difference in either the fln/+, fln/fln neurons with respect to the MMP values in comparison to +/+ (Fig. 7D). We further labeled mitochondria with Mito Tracker to measure the mitochondrial activity, morphology and mobility. As shown in Fig. 7E, the shape and size of mitochondria in the fln/+, fln/fln neurons did not differ significantly from the +/+ (Fig. 7E, F). We also quantitated mitochondrial movement within neurites of cortical neurons. Representative kymographs for each genotype were shown in Fig. 7G. Our results have demonstrated that the mutant neurons, both the fln/+ and fln/fln, did not exhibit significant difference in the moving speed (Anterograde, Fig. 7H; Retrograde, Fig. 7I), in the percentile of mitochondria moving in the retrograde (Fig. 7J) or total moving mitochondria (Fig. 7K). We also collected brain tissues and performed Western analysis (Fig. 7L). Our results demonstrated that fln/+ and fln/fln samples did not differ significantly from the +/+ in the protein level for either Drp1 (Fig. 7M) or TOM20 (Fig. 7N), a mitochondrial outer membrane protein^86, 87^. Taken together, our data have demonstrated that mitochondrial morphology and function in E18 CNS neurons is not markedly affected in the mutant CMT2B mouse model.

**Figure 7.**
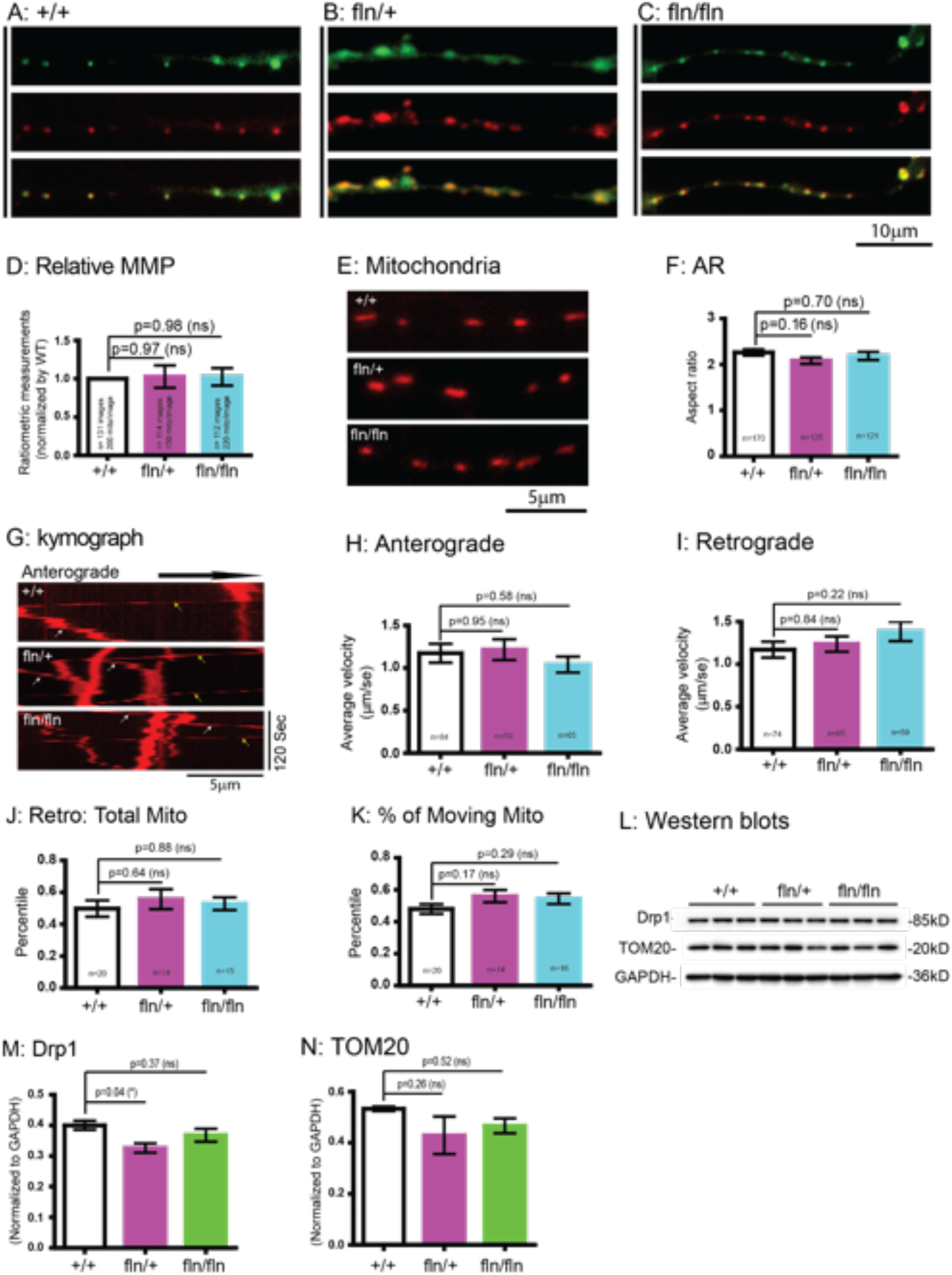
Analysis of mitochondria in E18 cortical neurons from CMT2B mutant mice. E18 cortical neurons from +/+, fln/+ fln/fln embryos were dissected, cultured and assayed for mitochondrial MMP, motility as in Fig. 6. A-C: Representative images of mitochondria in neurites of E18 cortical neurons of +/+, fln/+ and fln/fln. D: Ratiometric measurements of MMP. In E, Representative images of mitochondrial morphology in cortical neurons of +/+, fln/+, fln/fln. F: Aspect ratio measurements. Mitochondrial transport in neurites of cortical neurons of +/+, fln/+, flnfln was also captured by time lapsed imaging. The image series were used to generate kymographs and a representative image for each genotype is shown in G. White arrows indicate trafficking in anterograde direction, yellow arrows indicate trafficking in retrograde direction. Average velocities of axonal mitochondria moving in the anterograde (H) and retrograde (I) direction. J: % of retrogradely moving mitochondria. K: % of total moving mitochondria. Results are shown as mean ± SEM. Significance analysis was carried out using Prism. Statistical significances were calculated by One-Way ANOVA. n.s.= non significance. All *p* values are shown in the graphs. In L, 2 month-old brain lysates from +/+ (n=3), fln/+ (n=3), fln/fln (n=3) were generated and 20 μg proteins were analyzed by SDS-PAGE and immunoblotting with indicated antibodies. The relative levels of Drp1 (M) and TOM20 (N) were quantitated and normalized against GAPDH using BioRad-Image Lab. Results are shown as mean ± SEM. Significance analysis was carried out using Prism. Statistical significances were calculated by One-Way ANOVA. n.s.= non significance. All *p* values are shown in the graphs.

## Discussion

Increasing evidence has suggested an important role of Rab7 in regulating mitochondrial morphology and function^64–66, 88^. Using fibroblasts from human CMT2B patients harboring Rab7^V162M^ mutation as well as a mouse model with knockin expression of Rab7^V162M^, we have demonstrated significant mitochondrial fragmentation in fibroblasts of both human CMT2B patients and the knockin mouse model. Mitochondria also exhibited similar phenotypes in mouse primary sensory DRG neurons of the knockin mouse model. Our study suggests that the effects are mediated by both the increase in GTP binding of Rab7^V162M^ mutant and enhanced Drp1 activity.

Mitochondrial size is regulated by both fusion and fission. Rab7 has been found to influence both Drp1 and mitofusin (MFN). Dysregulation of either Drp1 or MFN or both will affect the dynamics of mitochondrial morphology and function. Excessive mitochondrial fragmentation foretells both structural and functional degeneration of the peripheral neurons. Recent studies have shown that Rab7 was co-immuno-precipitated with MFN2, the mitochondrial fusion protein^11^; Dominant inhibitory mutation(s) in MFN2 is associated with CMT2A^17, 19, 89, 90^. CMT2A also manifests significant mitochondrial fragmentation and axonal transport defects^16, 17, 19, 91^. Targeting mitochondrial fragmentation provides a promising therapeutic strategy for treating CMT2 diseases as demonstrated elegantly by a recent study^18^. Intermittent activation of endogenous mitofusin with a small molecule normalized CMT2A neuromuscular dysfunction, reversed pre-treatment axon and skeletal myocyte atrophy, and enhanced axon regrowth by increasing mitochondrial transport within peripheral axons and promoting in vivo mitochondrial localization to neuromuscular junctional synapses^18^. It will be interesting to test if a similar strategy will be effective in treating CMT2B in future studies.

Rab7 was also found to regulate phosphorylation of Drp1^60^ and siRNA-mediated knocking down of endogenous Rab7 decreased phosphorylation of Drp1 (pS616). It has been demonstrated that phosphorylation of Drp1 (pS616) is required for Drp1 to promote mitochondrial fission^61–63^. In our present study, treating either MEFs or E18 cultured DRG neurons of CMT2B mouse model with Mdivi-1 normalized mitochondrial aspect ratio. These data have provided evidence that phosphorylation of Drp1 (pS616) is likely upregulated by increased GTP binding to Rab7^V162M^, leading to mitochondrial fragmentation.

Our results differ from previous reports that expression of CMT2B Rab7 mutations increased the mitochondrial length in axons of frog RGCs^66^ and expression of the constitutive active Rab7 mutant (Rab7^Q79L^) inhibited mitochondrial fission in HeLa cells^64^. Even in primary hippocampal and cortical neurons cultured from E18 embryos of our CMT2B Rab7^V162M^ mutant mice (Fig. 6, 7), we did not detect a significant change in mitochondrial size as compared to wt E18 neurons. One potential explanation is that Rab7^GTP^, when expressed at physiological level, acts through phosphorylation of Drp1 to increase its activity. Expression of Rab7^V162M^ in both CMT2B patients and mouse model results in a modest increase in the level of Rab7^GTP^, which is sufficient to enhance Drp1 phosphorylation, leading to mitochondrial fragmentation. However, heterologous overexpression of Rab7 mutants may cause an overload of Drp1 by Rab7^GTP^, consequently, leading to inhibition, rather than activation, of Drp1.

Peripheral sensory neurons are particularly afflicted in CMT2B. Consistent with these clinical observations, our data have shown that mitochondrial morphology and axonal transport were dysregulated in E18 primary cultured DRG neurons, but not in E18 primary hippocampal and cortical neurons. This is both interesting and intriguing at the same time, given that Rab7 is ubiquitously expressed. One likely explanation is the selective expression of peripherin, an intermediate filament protein expressed primarily in peripheral neurons. Rab7 has been shown to interact with peripherin and the disease-causing Rab7 mutant proteins all exhibited increased propensity for binding to peripherin. As such, peripherin is dissociated from neuronal filaments leading to the destabilization of axonal structure^92^. This will significantly impact axonal function and perhaps the mitochondrial morphology and function, particularly in neurons with long axons.

One of the surprising findings in our current study is the observation that both the fibroblasts from human CMT2B patients and the MEFs from CMT2B mouse model showed significant mitochondrial fragmentation. Although the exact reason(s) is currently not known, we speculate that mitochondrial dysfunction in fibroblasts may impair the ability for skin wound repair, contributing to the fact that CMT2B patients are more prone to skin ulcers. Interestingly, we have recently demonstrated that cell migration was significantly altered in human CMT2B patient Rab7^V162M^ fibroblasts ^93^, that may indicate changes in the ability for skin wound repair in CMT2B.

In conclusion, our current study has established that Rab7 plays an important role in regulating mitochondrial morphology and function. Additionally, Rab7 is involved in many other important facets of cellular processes from endocytic trafficking, axonal transport, autophagy, to lysosomal function. Exactly how these processes are impacted by CMT2B related Rab7 mutations warrant future investigations.

## Materials and Methods

### Ethical Statement

All experiments involving the use of laboratory animals have been approved by the Institutional Animal Care and Use Committee of University of California San Diego. Surgical and animal procedures were carried out strictly following the NIH Guide for the Care and Use of Laboratory Animals.

### Reagents and antibodies

10x Hanks’ Balanced Salt solution (HBSS), 2.5% Trypsin (10x), 100x Penicillin-Streptomycin (P/S), 100x GlutaMax, 50x B27, Neuronal basal media were all purchased from Invitrogen. DNase I grade II from bovine pancreas (Roche, Cat#10104159001) was dissolved in 1x HBSS at 10 mg/ml (10x) and filter-sterilized. Collagenase was from Advanced BioMatrix (#5030). MEM containing GlutaMAX was from Thermal Fisher. DMEM containing high glucose (4.5g/L) was from ThermoFisher. FBS was from Mediatech Inc (Cat# 35-010-CV). 0.1% Poly-L-Lysine (Cultrex® Poly-L-Lysine) was from Trevigen (Gaithersburg, MD; Cat# 3438-100-01). Mouse NGF was purified from submaxillary glands as described previously^94^. All other chemicals and lab wares were from Bio-Rad, Fisher, Sigma, VWR.

Mito Tracker™ Red FM (ThermoFisher, Cat#M22425) and JC-9 Dye (Mitochondrial Membrane Potential Probe) (ThermoFisher Cat# D22421) were purchased from Invitrogen (Carlsbad, CA). Mdivi-1 was purchased from MedChemExpress (Cat# HY-15886). CID11067700 was from Sigma (cat# SML0545). Both Mdivi-1 and CID11067700 were dissolved in dimethyl sulfoxide solution (DMSO from Sigma). Rabbit monoclonal Ab against Drp1 (D6C7) was from Cell Signaling Technology (CST#8570S), rabbit anti-TOM20 (sc-11415) and rabbit anti-GAPDH (sc-47724) IgGs were from Santa Cruz Biotechnology. Goat anti-rabbit IgG-HRP conjugates were purchased from Jackson ImmuoResearch Laboratories. All antibodies were used at dilutions following manufacture instructions.

### Human fibroblasts from healthy control and CMT2B^V162M^ patient

Human fibroblasts were isolated from skin of patients affected by CMT2B (two 46 and 53- years old men and 52-year old woman) and from healthy control individuals age and sex matched^33, 47^. These CMT2B patients are from the first identified Italian family affected by this pathology, which was diagnosed at Azienda Ospedaliera Universitaria “Federico II” of Naples. Clinical history and symptoms of the CMT2B patients have been previously described^33^. Informed consent was obtained in compliance of the Helsinki Declaration. The Study was approved by the local Ethics Committee (Ethical Committee Approval Protocol # 107/05). All samples were anonymously encoded to protect patient confidentiality. Tissue digestion and cell isolation were performed as previously described^47^.

Fibroblasts were grown in high-glucose Dulbecco’s modified Eagle’s medium (DMEM, Corning, NY, USA) supplemented with 10% (v/v) fetal bovine serum (FBS), 1% (v/v) L-glutamine, 1% (v/v) penicillin/streptomycin (Sigma-Aldrich, St. Louis, MO, USA) at 37 ◦C in a humidified atmosphere of 5% CO2. For mitochondrial labeling with Mitotracker, cells were seeded on coverslips placed in 12-well plates as described in the Measurement of mitochondrial morphology and movement section.

### Rab7^V162M^ knockin mouse model

We have made a knockin mouse model (C57BL6) for CMT2B by changing 484G to A (V162>M) in Rab7 Exon 5 (Supplemental Fig 1A). The mutated allele was introduced into the mouse genome, together with the selection marker NeoR and two LoxP sites. The LoxP sites will facilitate studies in which selective deletion of the mutated allele is needed using Cre. Genotyping was performed using the PCR primer pair (Supplemental Fig 1B). The wild type (wt) gives rise to a 373 bp fragment while the mutant allele (Rab7V162M:Neo) produces a 490 bp. We obtained all three genotypes: wt (+/+), heterozygote (fln/+) and homozygote (fln/fln) with a typical Mendelian segregation ratio. Both the fln/+ and fln/fln pubs survive to full adulthood.

### Mouse embryonic fibroblasts (MEFs) culture and maintenance

Abdominal skin tissues were excised from wild +/+, fln/+ and fln/fln E18 mouse embryos and were cut with scissors into about 1×1mm^2^ small pieces. Following quick rinse in HBSS with 1% P/S, the skin tissues were dissociated in 0.25% Trypsin with 1000 U/ml collagenase in HBSS at 37°C for 30 mins. And then DNase I (1 mg/ml, final concentration) was added into the digestion. Dissociated fibroblasts were centrifuged and cultured in medium (High glucose DMEM with 15% FBS, 1% P/S). Half of the medium was replaced the following day and then every other day until conclusion of the experiments.

### Dorsal root ganglion (DRG) neuronal culture and maintenance

Established protocols were followed to set up DRG neurons from E18 mouse embryos of +/+, fln/+ and fln/fln ^95^. Briefly, DRG tissues from mouse E18 embryos were extracted from both sides of L2, L3, L4, L5, L6 levels and extensively rinsed in HBSS with 1% P/S, followed by dissociation in 0.25% trypsin at 37°C for 20 mins. DRG neurons were triturated with fire-polished glass pipets and dissociated DRG neurons were cultured in plating media (MEM containing GlutaMAX with 100ng/ml NGF, 10% FBS). The culture dishes were pre-coated with poly-L-lysine (Invitrogen). Plating medium was completely replaced with maintenance medium (MEM containing GlutaMAX with 100ng/ml NGF, 1% FBS). Arabinosylcytosine (AraC) was added at a final concentration 1µM into maintenance medium to suppress the proliferation of non-neuronal cells like fibroblasts, Schwann cells the following day. Regular maintenance medium and selection maintenance medium containing AraC were used alternately every other day until the conclusion of the experiments.

### Cortical and hippocampal neuronal culture and maintenance

Cortical and hippocampal neuronal cultures were carried out following published protocols^96^. Briefly, cortical or hippocampal tissues were dissected from +/+, fln/+, fln/fln E18 mouse embryos and extensively rinsed in HBSS with 1% P/S, followed by dissociation in 0.25% trypsin with 1 mg/ml DNase I. Neurons were isolated and plated with plating media (Neurobasal media with 10% FBS, 1xB27,1xGlutaMAX) onto glass coverslips at appropriate density. Both the coverglasses and plates were pre-coated with 0.1% poly-L-lysine (Invitrogen). Plating medium was replaced with maintenance medium (Neurobasal media, 1xB27, 1xGlutaMAX) the following day. Only 2/3 of the media was replaced every other day until the conclusion of the experiments.

### Measurement of mitochondrial morphology and movement

Mito Tracker™ Red FM was diluted into minimal essential media (MEM). MEFs were treated with a final concentration of 50 nM of Mito Tracker™ Red FM at 37°C for 30 mins. After quick rinse with MEM, the cultures were live-imaged under a 63X objective lens with a zoom factor of 1.6X using a Leica DMi8 Live Imaging Microscope. The analysis and quantification of mitochondrial morphology and network complexity were carried out as described in the section of **Quantification of mitochondrial size, network complexity** according to the published methodologies^73, 74^.

DRG neuronal cultures were treated with a final concentration of 100nM of Mito Tracker™ Red FM at 37°C for 40 mins. Cortical and hippocampal neurons were treated with a final concentration of 50nM of Mito Tracker at 37°C for 30 mins. Cultures were then quickly rinsed and imaged to track mitochondrial trafficking. Time-lapsed image series were captured at 1.0 frame/2 sec for a total 2 mins. For DRG cultures, the images were taken under a 63X objective lens. For cortical and hippocampal neurons, the images were taken under a 63X objective lens with a zoom 1.6X. Kymographs were generated, and analysis and quantification of mitochondrial transport were carried out as described previously^96^. The width and height of individual mitochondrion were measured, and the aspect ratio as a measurement for mitochondria size.

Human fibroblasts were seeded into microscopy chambers (8 well μ-slide, Ibidi GmBh, Martinsried, Germany) and, after 24 h, incubated with 50 nM MitoTracker Red CM-H2XROS (ThermoFisher Scientific) for 40 min at 37 ◦C in DMEM medium without serum. After 3 washes in PBS, L-15 medium was added, and cells were imaged by confocal microscopy. Fluorescence images were captured using a confocal laser scanning microscope (CLSM) (Zeiss, LSM 700, Germany) equipped with a laser diode emitting at 405 nm, an argon-ion laser for excitation at 488 nm, and a helium-neon laser for excitation at 555 nm. Images were taken with a Plan-Apochromat 63.0 × 1.40 oil-immersion objective DIC M27. The images were acquired using ZEN Black Edition 2011 software (Zeiss, Jena, Germany).

### Ratiometric measurement of mitochondrial membrane potential

JC9 is a cationic dye which binds to mitochondria and emits green fluorescence (∼525 nm) independent of mitochondrial membrane potential (MMP). In the case of mitochondrial membrane hyperpolarization, JC9 aggregates and gives off red fluorescence (∼590 nm). Therefore, the intensity ratio of red: green can be used to measure the MMP and indicates the mitochondrial activity^97^.

Primary cortical and hippocampal neurons were loaded with maintenance media containing JC-9 dye (final concentration: 1μg/ml). After incubation in 37°C for 25 min, neurons were live-imaged under a 63X objective lens with a zoom factor of 1.6X. Both green and red channel images were captured. NIH Image J (Fiji) software was used to measure the intensity of green and red fluorescent signals. And the ratio of red/ green fluorescent intensity was calculated.

### Treatments with Mdivi-1 and CID1067700

MEFs were treated with Mdivi-1 at a final concentration of 50µM in high-glucose DMEM. Mdivi-1 at 50µM has been shown previously effective in inhibition of Drp1 activity in vitro^82, 98^.

After incubation at 37°C for 30 mins, fln/fln fibroblasts were treated with Mito Tracker™ Red FM and imaged lived as described above.

CID1067700 was diluted into high-glucose DMEM and applied to fln/fln MEF cultures at the final concentration of 1µM and 10µM. After incubation at 37°C for 30 mins, fibroblasts were treated with Mito Tracker™ Red FM and were live-imaged. DMSO (0.1%, final concentration) was applied into medium as control treatment.

For fln/+ DRG cultures treatment, Mdivi-1 was diluted into DRG maintenance media at the final concentration of 50µM, 100µM, 200µM for 1 hr at DIV5, followed by Mito Tracker™ Red FM treatment. Live-imaging was carried out to measure the aspect ratio of mitochondria as described above to define the maximum rescuing effect of Mdivi-1 on mitochondrial fragmentation. For rescuing analysis, fln/fln DRG neurons were pre-incubated in maintenance medium containing Mdivi-1 (100µM, final concentration) for 1 hr, followed by incubation with Mito Tracker™ Red FM. DMSO-treated fln/fln DRG sensory neurons were as control cultures.

### Quantification of mitochondrial size, network complexity

Established methods for multidimensional measurements of mitochondrial morphology was used to quantitate the mitochondrial size and network complexity^73, 74^. The Mitochondria Analyzer plugin can be downloaded from http://sites.imagej.net/ACMito/ and installed in Fiji of NIH ImageJ. Briefly, healthy mitochondria are generally mobile and tubular in shape and exist in complex networks, whereas cells undergoing profound stress or entering apoptosis often display swollen and fragmented mitochondria. The mitochondria analyzer is designed to measure both the size and shape of individual mitochondrion, as well as network complexity; The mitochondrial size can be measured in area, perimeter. The shape is reflected by the aspect ratio (maximal:minimal diameter) and by the form factor. The complexity of mitochondrial network is revealed by the number of branch joints, branches, and branch length.

Follow the instruction^74^, we defined the mitochondrial morphology to three different levels of complexity: Level 1 (L1) was for the highly complex mitochondrial network with little or no fragmented mitochondria; Level 2 (L2) represented less complex mitochondrial network with fragmented mitochondria; and Level 3 (L3) was for highly fragmented mitochondria with little or no network. For 2D analysis, the image was first processed, and a threshold was applied; The blocking size of 1.05/1.25/1.45μm and C-value of 5/9/13 were applied for L1, L2 and L3, respectively. The resulting binary image was used as the input for the “Analyze Particles” command (size = 100/70/40 pixels -infinity, circularity = 0.00–1.00, for level 1/2/3). Note that only mitochondria larger than 100/70/40-pixel units were taken into consideration in those 3 conditions, respectively. The output measurements generated for “area” and “perimeter” represented the average size of mitochondria; The shape of mitochondria was revealed by the values for both the aspect ratio (AR) and the Form Factor (FF). FF was derived as the inverse of the “circularity” output value to take into consideration of curvatures.

For network connectivity analysis, the “skeletonize 2D/3D” command was applied to the image following application of the threshold to produce a skeleton map, and the “Analyze Skeleton” command was used to calculate the numbers of branches and branch junctions. Total branch length and average branch length were also generated from the skeletonized network^74^.

### Western blot analysis of the expression of mitochondria-related proteins

The whole brains were dissected from +/+, fln/+, fln/fln mice at 2 months of age. The brains were homogenized in HB (0.32 M Sucrose, 10mM HEPES pH7.5, 0.2mM phenylmethylsulfonyl fluoride), followed by centrifuging at 1000 rpm at 4°C for 5 mins. The resulting supernatants were collected and measured to decide the concentration of total proteins. Equal amounts of proteins were denatured and separated by 12.5% of SDS-PAGE (sodium dodecyl sulfate polyacrylamide gels). Separated proteins were transferred to PVDF membranes, blocked in 5% nonfat milk in TBST (150 mM NaCl, 10 mM Tris-HCl, pH 7.5 and 0.1% Tween 20). Immunoblotting was performed with anti-Drp1 antibody (1/1000), anti-TOM20 antibody (1/2000), anti-GAPDH antibody (1/10,000), and then with corresponding secondary antibodies (goat anti-rabbit, 1:10,000). The blots were developed in ECL-Clarity (BioRad) and were imaged using ChemiDoc XRS+ (Bio- Rad). The blots within linear exposure ranges were quantitated using the ImageLab 6.0.1 software (BioRad).

---

**Supplemental** **Fig 1. A CMT2B Rab7^V162M^ knockin mouse model (C57BL6).**

A: a schematic shows the knockin strategy. B: PCR primers and tail DNA genotyping results of wt (+/+), heterozygote (fln/+) and homozygote (fln/fln). C: both fln/+, fln/fln mice survived to adulthood.

**Supplemental** **Fig 2. Mitochondrial morphology in soma of DRG sensory neurons**

A: Representative images of Mito Tracker in E18 DRG sensory neurons from +/+, fln/+, fln/fln embryos. B: The phase contrast images corresponding to A.

**Supplemental** **Fig 3. Mitochondrial morphology in soma of hippocampal neurons**

A: Representative images of Mito Tracker in E18 hippocampal neurons from +/+, fln/+, fln/fln embryos. B: The phase contrast images corresponding to A.

**Supplemental** **Fig 4. Mitochondrial morphology in soma of cortical neurons**

A: Representative images of Mito Tracker in E18 cortical neurons from +/+, fln/+, fln/fln embryos. B: The phase contrast images corresponding to A.

**Supplemental Movie S1**. Axonal trafficking of mitochondria in DRG sensory neurons from +/+ E18 embryos

**Supplemental Movie S2.** Axonal trafficking of mitochondria in DRG sensory neurons from fln/+ E18 embryos

**Supplemental Movie S3.** Axonal trafficking of mitochondria in DRG sensory neurons from fln/fln E18 embryos

## Acknowledgement

We thank Ms Pauline Yue Hu for technical support.

## Author contributions

YG performed the majority of the experiments, generated the data, and helped with writing the manuscript; FG generated the data from human fibroblasts; MH was responsible for performing and quantitating mitochondrial size and complexity; AP provided help in collecting raw data; KS, WY generated the knockin mouse data (Supplemental Fig 1); SJ, TS, TG, OD, LS helped to quantitate mitochondrial size and axonal transport data; YZ, JD provided the knockin mouse model; CB, CW conceived the experimental design and wrote the manuscript.

