## Supplementary figures and images for "Mitochondria dysfunction in Charcot Marie Tooth 2B Peripheral Sensory Neuropathy"

### supplemental figures and movies

#### Slide 1
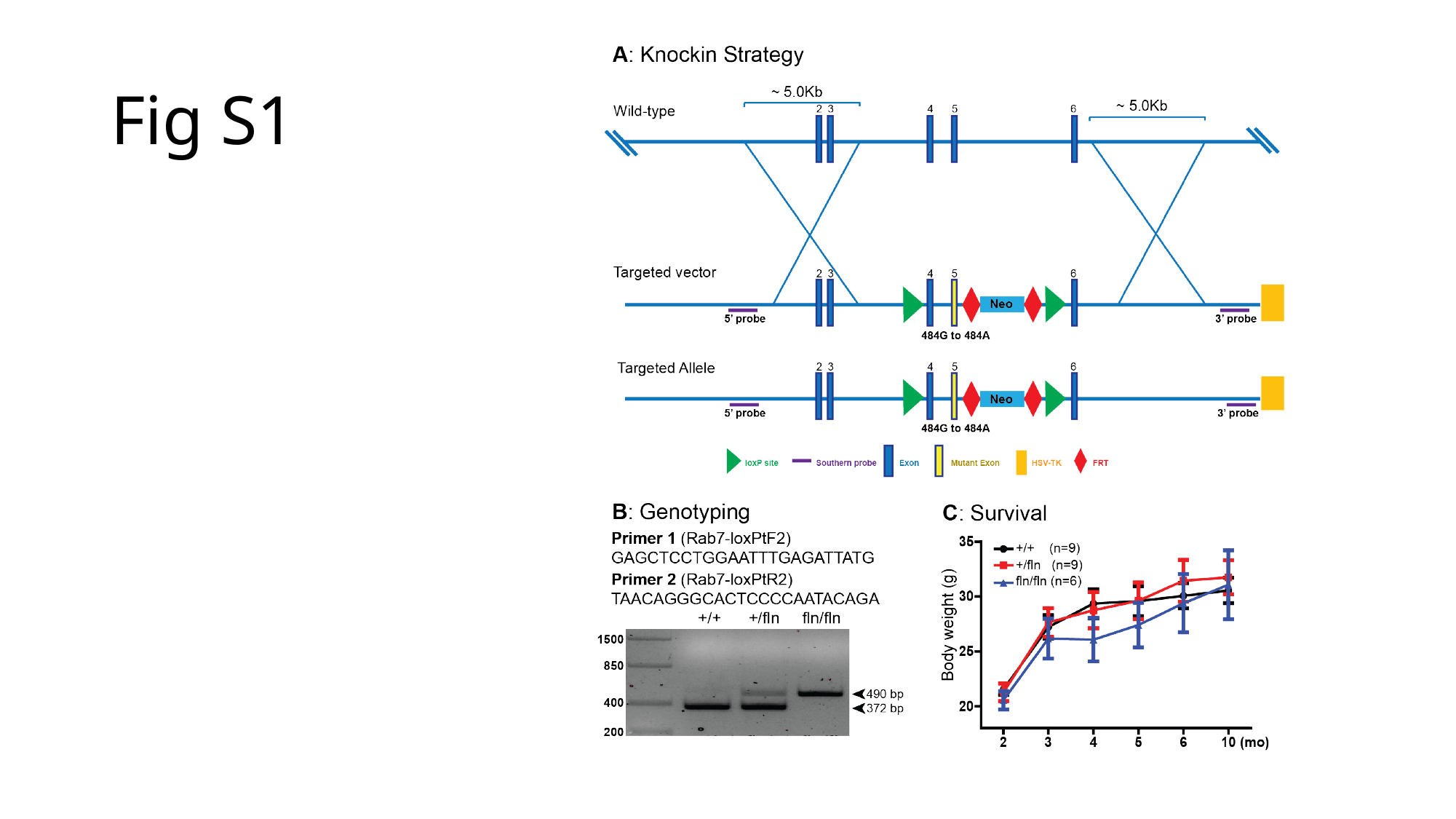

##### Fig S1

#### Slide 2
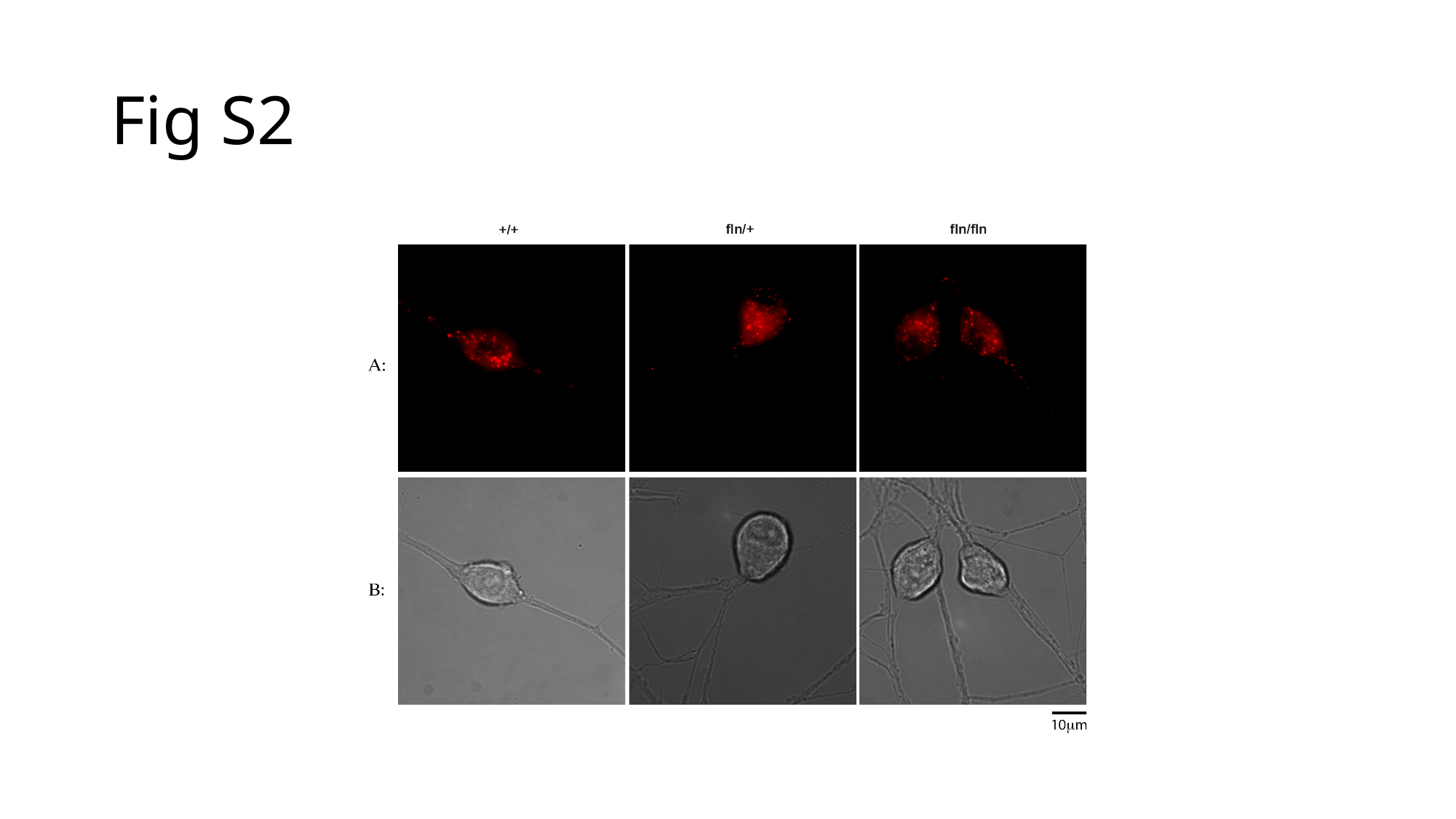

##### Fig S2

#### Slide 3
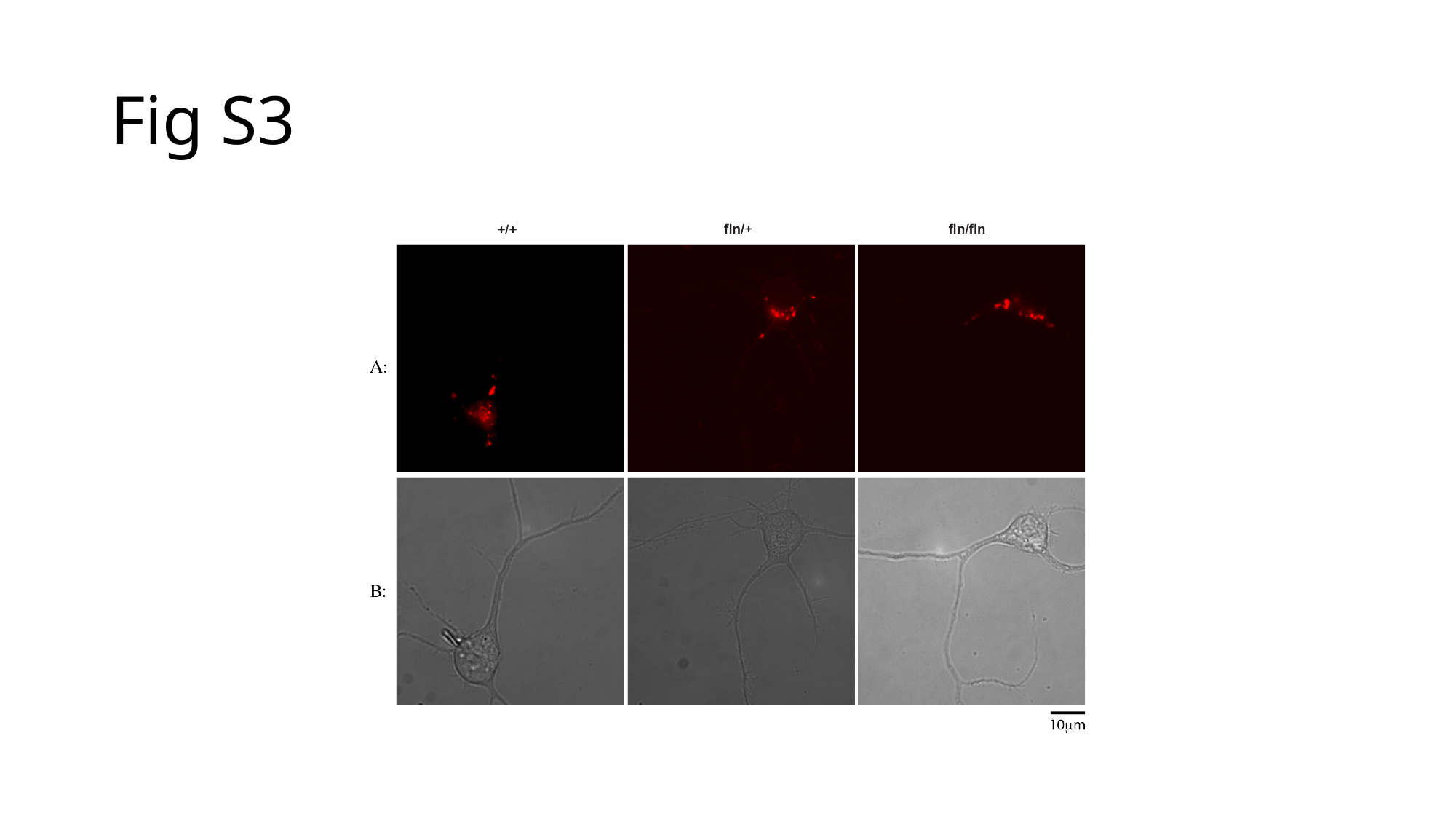

##### Fig S3

#### Slide 4
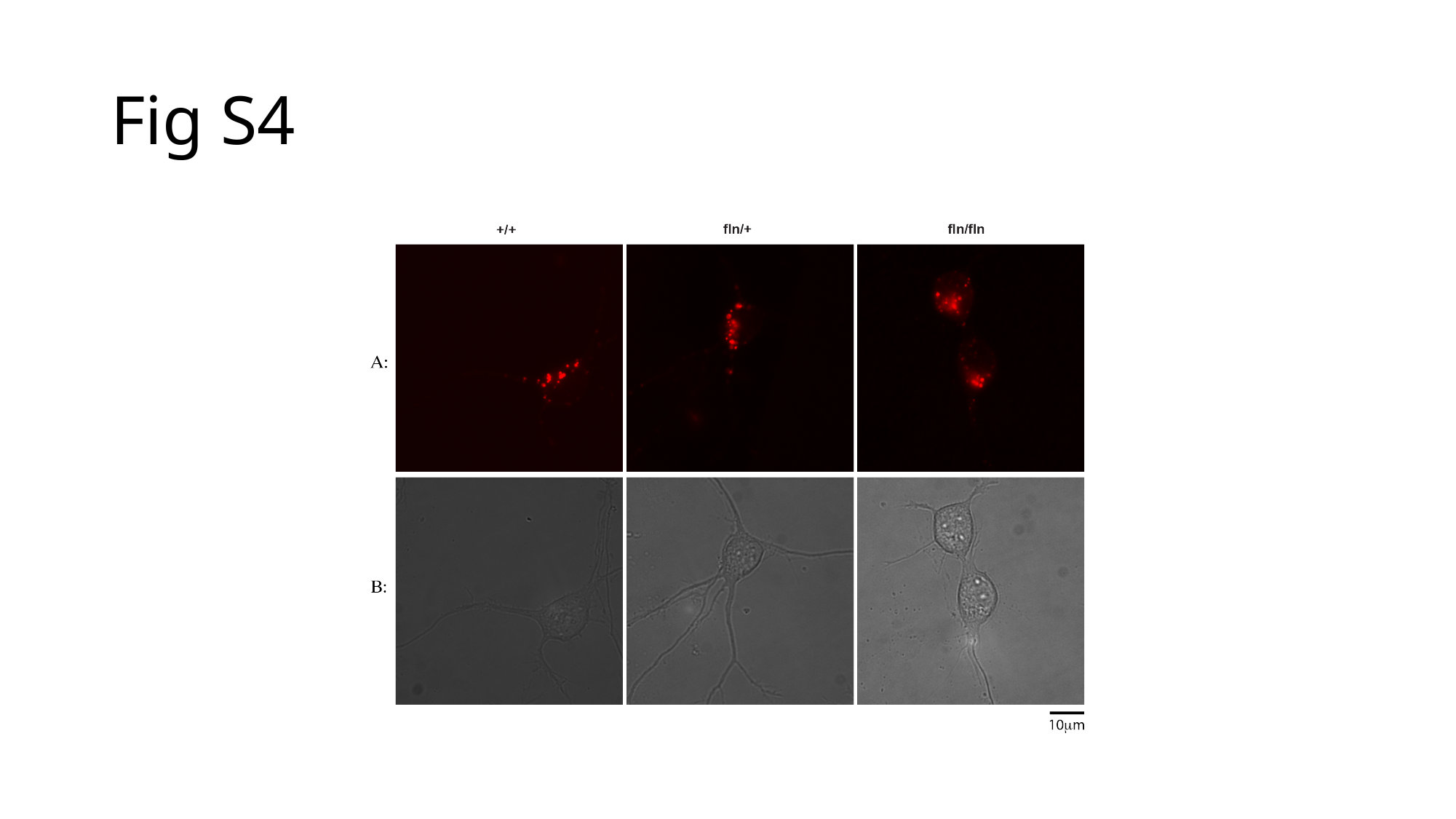

##### Fig S4

#### Slide 5
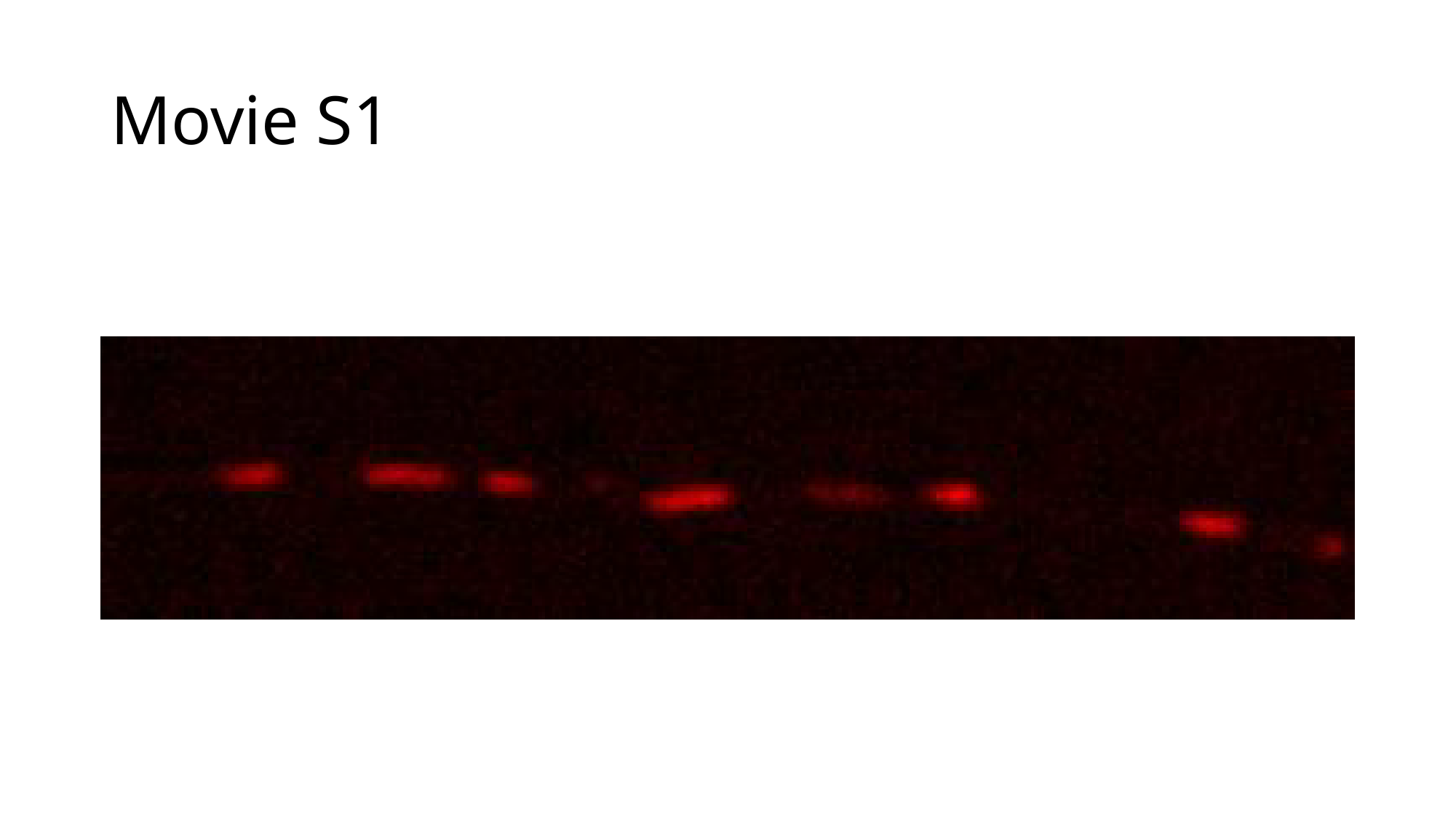

##### Movie S1

#### Slide 6
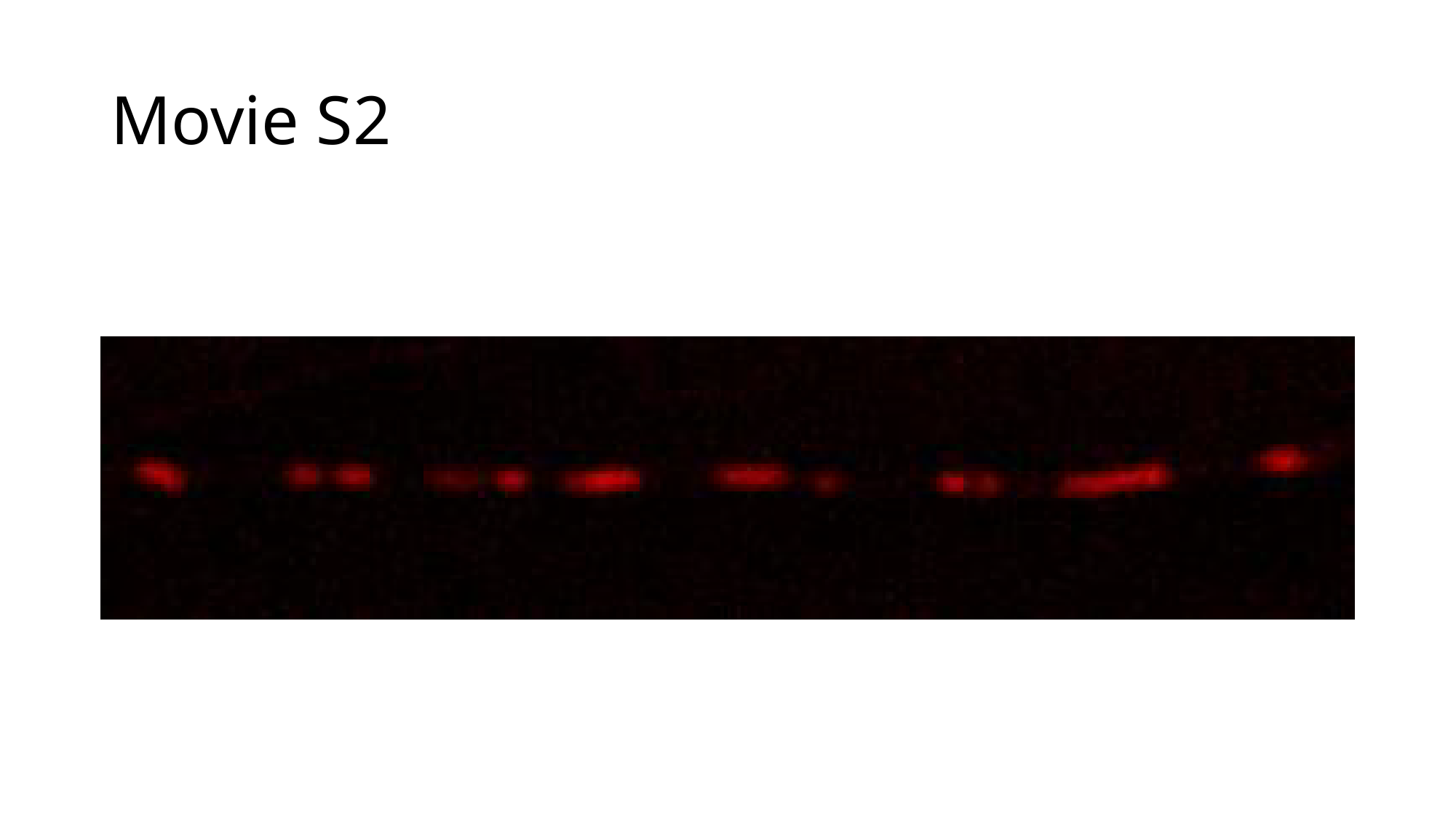

##### Movie S2

#### Slide 7
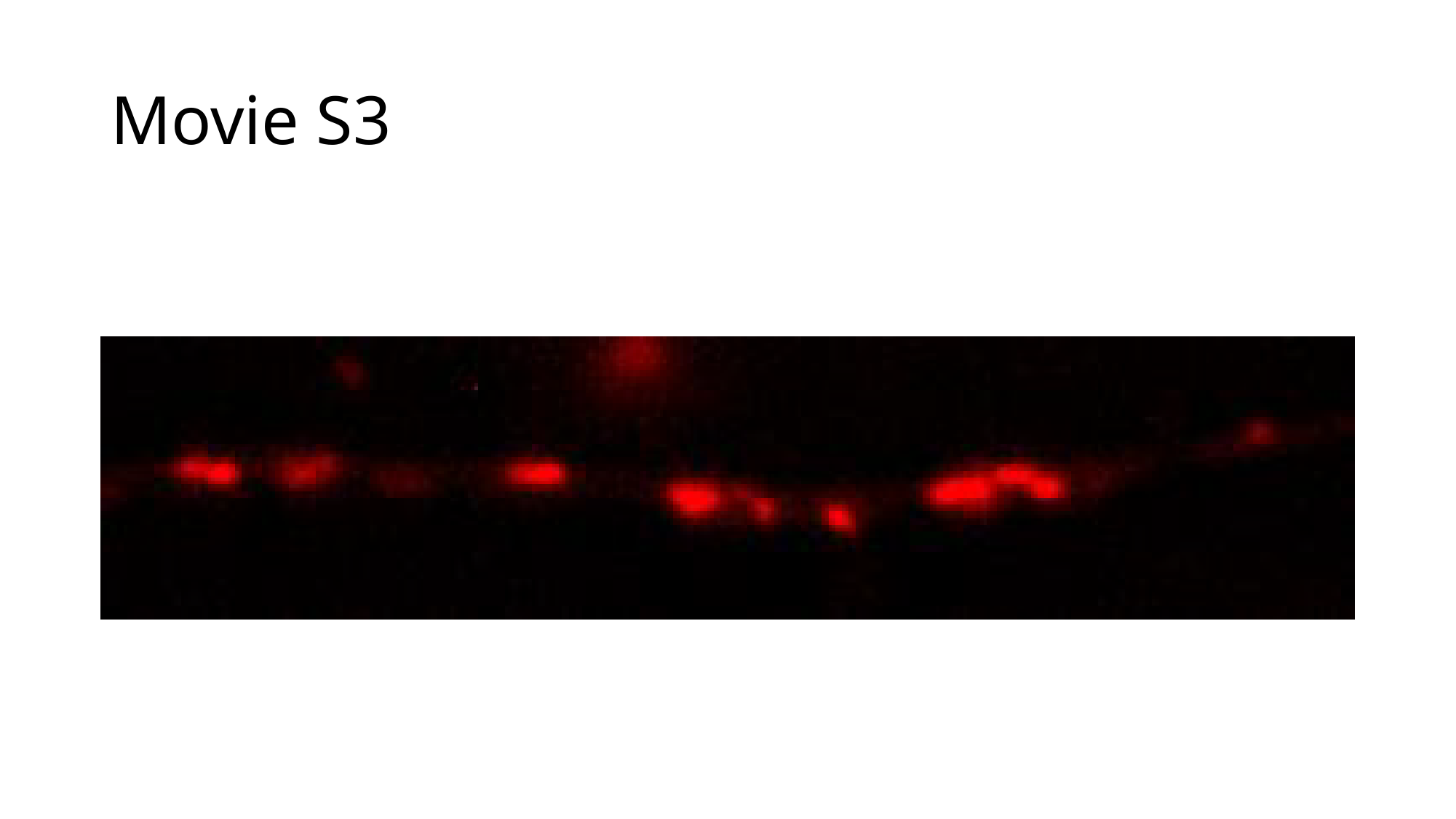

##### Movie S3
